# Orchestration of alternative splicing regulates bone marrow mesenchymal stem cells fate during aging

**DOI:** 10.1101/2022.05.27.493685

**Authors:** Ye Xiao, Guang-Ping Cai, Xu Feng, Qi Guo, Yan Huang, Tian Su, Chang-Jun Li, Xiang-Hang Luo, Yong-Jun Zheng, Mi Yang

## Abstract

Senescence and change of differentiation direction in bone marrow stromal cells (BMSCs) are two of the most important causes of age-related bone loss. As an important post-transcriptional regulatory pathway, alternative splicing (AS) regulates diversity of gene expression. However, the role of AS in BMSCs during aging remains poorly defined. Here we identify AS in specific genes disrupt gene expression pattern and result in age-related debility of BMSCs. We demonstrate the deficiency of splicing factor Y-box protein 1 (YBX1) result in mis-splicing in genes such as *Fn1, Taz, Sirt2* and *Sp7*, further contributing to senescence and shift in differentiation direction of BMSCs during aging. Deletion or over-expression of YBX1 in BMSCs accelerate bone loss or stimulate bone formation in mice. Notably, we identify a small compound sciadopitysin which attenuate the degradation of YBX1 and attenuate bone loss in old mice. Our study demonstrates elaborately controlled RNA splicing governs cell fate of BMSCs and provides a potential therapeutic target for age-related osteoporosis.

**Summary:** This study demonstrates that YBX1 deficiency induces pre-mRNA mis-splicing and causes senescence and shift in differentiation direction of BMSCs and further accelerates aging-related bone loss. This study identifies Sciadopitysin could reverse this process by targeting YBX1.

## Introduction

Alternative splicing of precursor mRNA (pre-mRNA) is an important post-transcriptional regulatory pathway of more than 90% of multiexon protein-coding genes in mammals, enabling cells to generate abundant transcript and protein diversity from a limited number of genes^1^. Most of human genes undergo alternative splicing ^2^ in a variety of physiological and pathological process such as mesenchymal stem cell differentiation, tissue and organ development, aging and tumorigenesis ^3–7^. Dysregulation of pre-mRNA splicing is associated with aging and a large proportion of age-related changes in alternative splicing are associated with alternations of the expression of splicing factors ^8–10^.

Bone marrow stromal cells (BMSCs) have the ability to differentiate into a variety of cell types including osteoblasts, adipocytes and chondrocytes ^11–13^. With age, BMSCs become more inclined to differentiate into adipocytes rather than osteoblasts, resulting in bone marrow fat accumulation and bone loss^14–17^. However, the molecular mechanism which regulates age-associated BMSCs lineage shift remains elusive. In BMSCs, the expression of genes related to senescence such as *P53/P16*, or key transcriptional factors related to differentiation such as *Runx2* or *Ppargγ* could be regulated by alternative splicing ^18–21^. Changes of splicing factors and variable of AS events in BMSCs genes may be critical in BMSCs senescence and fate determination during aging.

Y-box binding protein 1 (YBX1) is a multifunctional protein known to participate in a wide variety of DNA/RNA-dependent events including DNA reparation and transcription, pre-mRNA splicing, mRNA stability and translation ^22–24^. YBX1 has been reported to control the expression of pluripotency-related genes in embryonic stem cells ^25^. YBX1 also could regulate multiple biological activities including cell proliferation, differentiation, senescence, apoptosis, and tumor development ^26–29^. It has been reported that YBX1 restrained cellular senescence by directly binds to the *p16^INK4A^* promoter and repressed the transcription of *P16^INK4A^* ^28^. In the past decades, as a splicing factor, the role of YBX1 in alternative splicing has been studied ^30–34^. Jayavelu AK et al reported that YBX1 mediated pre-mRNA splicing is the key mechanism of persistence of JAK2-mutated myeloproliferative neoplasms ^33^. Ma, S. et al. demonstrated that YBX1 decreased with aging in six tissues (bone marrow, brown adipose tissue, white adipose tissue, aorta, skin, liver) and participated in adipose stem cell maintenance in white adipose tissue ^35^. However, whether YBX1 regulates the fate of BMSCs via alternative splicing is unclear.

In the present study, we observed altered pre-mRNA splicing and changed gene expression pattern in BMSCs during aging and we further find that the expression of splicing factor YBX1 in BMSCs is decreased with aging in mice and human. YBX1 can stimulate osteogenic differentiation and restrain senescence of BMSCs by regulating a cluster of genes including *Fn1, TAZ, Sirt2* and *Sp7* as a splicing factor. Moreover, we discover a natural small compound, sciadopitysin, which can restrain the degradation of YBX1 and attenuate age-related bone loss. Our results demonstrate elaborately controlled RNA splicing regulates differentiation and senescence of BMSCs and suggest that YBX1 is a potential therapeutic target for age-related osteoporosis.

## Results

### 1. Dysregulated pre-mRNA alternative splicing and altered gene expression pattern in BMSCs during aging

Aging related decline in bone formation is closely associated with the debility of BMSCs. BMSCs isolated from 24-month-old mice showed significant higher level of senescence indicated by β-Gal staining and lower osteogenic differentiation potential indicated by Alizarin Red staining after osteogenic induction compared with BMSCs isolated from 2-month-old mice (Figure 1 A-C). Dysregulation of pre-mRNA splicing directly contribute to cell dysfunction and senescence. To investigate the splicing events and gene expression pattern in BMSCs during aging, we performed whole transcriptome resequencing and alternative splicing analysis in BMSCs isolated from 2-month-old and 24-month-old mice (Figure 1 D). We evaluated changes in pre-mRNA splicing by calculating “percentage spliced in”(ΔPSI) values of several major alternative splicing events with ΔPSI>0.1 and P value<0.05 (Figure 1 E). pre-mRNA splicing in BMSCs upon aging showed changes in alternative first exon event (35.98%), alternative 5’ splice site (15.09%), exon skipping (14.76%), intron retention (13.93%), alternative last exon (11.44%) and alternative 3’ splice site (8.46%) (Figure 1 E). Gene Ontology (GO) analysis showed that those alternative splicing events involved genes related with osteoblast differentiation and cellular senescence (Figure 1 F). Meanwhile, the BMSCs isolated from 24-month-old mice exhibit increased expression of a cluster of senescence and adipogenic differentiation related genes and decreased expression of a cluster of osteogenic differentiation related genes with at least a 2-fold change when compared with BMSCs isolated from 2-month-old mice (Figure 1 G). To investigate the potential splicing factor might be responsible to the altered splicing events in BMSCs during aging, we identified 92 RNA splicing proteins whose expression level changed with at least a 2-fold between BMSCs isolated from 2-month-old mice and 24-month-old mice, by combine analyzing RNA sequencing data with RNA binding proteins and RNA splicing proteins datasets (Figure 1 H, I). These differentially expressed splicing factors form a regulatory network whose functions are mainly enriched in terms related to various RNA metabolic processes (Figure 1 J). We further screen out 50 differentially expressed proteins with function enriched in mRNA splicing or regulation of RNA splicing (Figure 1 J, K)

**Figure 1.**
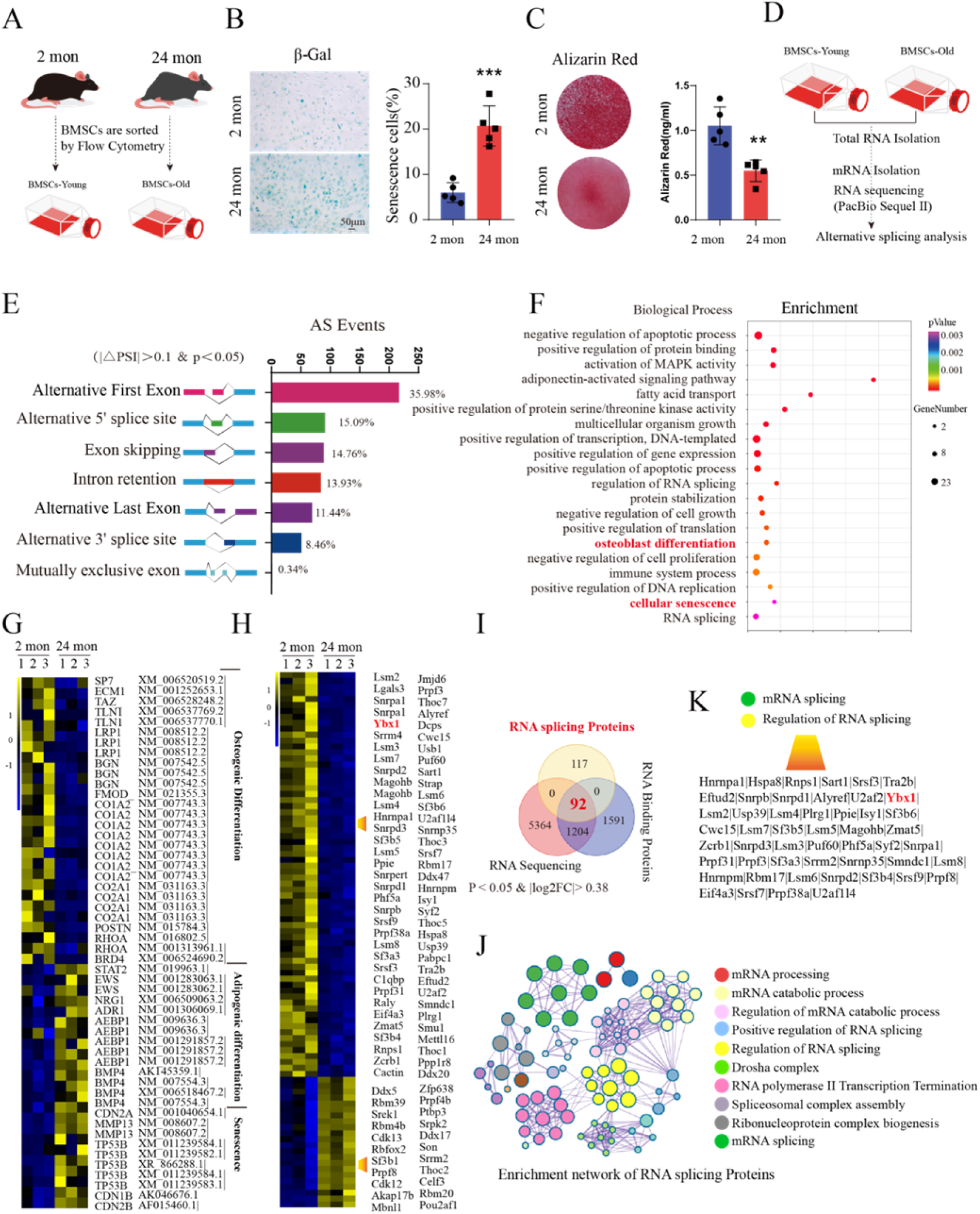
Dysregulated pre-mRNA alternative splicing and altered gene expression pattern in BMSCs during aging. (A) Schematic diagram of isolating and culture of BMSCs from 2-month-old and 24-month-old mice. (B) Representative images of β-Gal staining (left panel) and quantification of β-Gal positive cells (right panel) of BMSCs isolated from 2-month-old and 24-month-old mice. Scale bar: 50 μm. (C) Representative images of Alizarin Red staining at 21 days of osteogenic induction (left panel) and quantification of calcification (right panel) of BMSCs isolated from 2-month-old and 24-month-old mice by detecting the amount of Alizarin Red extracted from the matrix. (D) Schematic diagram of experimental process of alternative splicing analysis. (E) Histogram of the differentially spliced events between BMSCs isolated from 2-month-old and 24-month-old mice. (F) GO analysis of differentially expressed genes between BMSCs isolated from 2-month-old and 24-month-old mice. (G) Heat map of differentially expressed genes between BMSCs isolated from 2-month-old and 24-month-old mice. (H-I) Heat map of differentially expressed genes of RNA splicing proteins (H) and Venn diagrams of overlapping genes between differentially expressed genes, RNA binding proteins and RNA splicing proteins datasets (I) between BMSCs isolated from 2-month-old and 24-month-old mice. (J) Enrichment network representing the top 10 enriched terms of differentially expressed RNA splicing proteins between BMSCs isolated from 2-month-old and 24-month-old mice. Enriched terms with high similarity were clustered and rendered as a network, while each node represents an enriched term and is colored according to its cluster. Node size indicates the number of enriched genes, and the line thickness indicates the similarity score shared by two enriched terms. (K) The list of differentially expressed RNA splicing proteins between BMSCs isolated from 2-month-old and 24-month-old mice whose functions were clustered in mRNA splicing and regulation of RNA splicing. Data shown as mean ± SEM. **, *P* < 0.01; ***, *P* < 0.001; Student’s t test.

The changed splicing factors and pre-mRNA altered splicing events might play an important role in the functional debility of BMSCs during aging.

### 2. Splicing factor YBX1 regulated the fate decision and senescence of BMSCs and showed decreased expression during aging

Among these changed RNA splicing factors, YBX1, inactivation of which has been reported inducing apoptosis in mouse and primary human cells and cause regression of the malignant clones in vivo ^33^, displayed significantly lower expression in the BMSCs isolate from the older mice compared with that from young ones (Figure 1 H). We confirmed the lower expression level of YBX1 in BMSCs isolated from 24-month-old mice related to 2-month-old mice by immunofluorescence staining (Figure 2 A-C), western bolt (WB, Figure 2 D) and qPCR (Figure 2 E) analysis. As YBX1 performs its pre-mRNA splicing function mainly in nucleus, we further confirmed the lower YBX1 level in nucleus in BMSCs isolated from 24-month-old mice (Figure 2 D). The YBX1 level was also lower in cultured primary BMSCs from late passage (Supplemental Figure 1 A-C). Nestin positive cells in bone marrow represent a subset of messenchymal stem cell ^16,36^. Co-immunofluorescence staining of Nestin and YBX1 on the bone marrow confirmed the lower number of Nestin^+^ BMSCs and YBX1^+^cells in metaphysis area in femur of 24-month-old mice (Figure 2 B, C). Similarly, the YBX1 level in human BMSCs was also negatively correlated with age (Figure 2 F). These results indicated that slicing factor YBX1 might play roles in the altered splicing events in BMSCs during aging.

**Figure 2.**
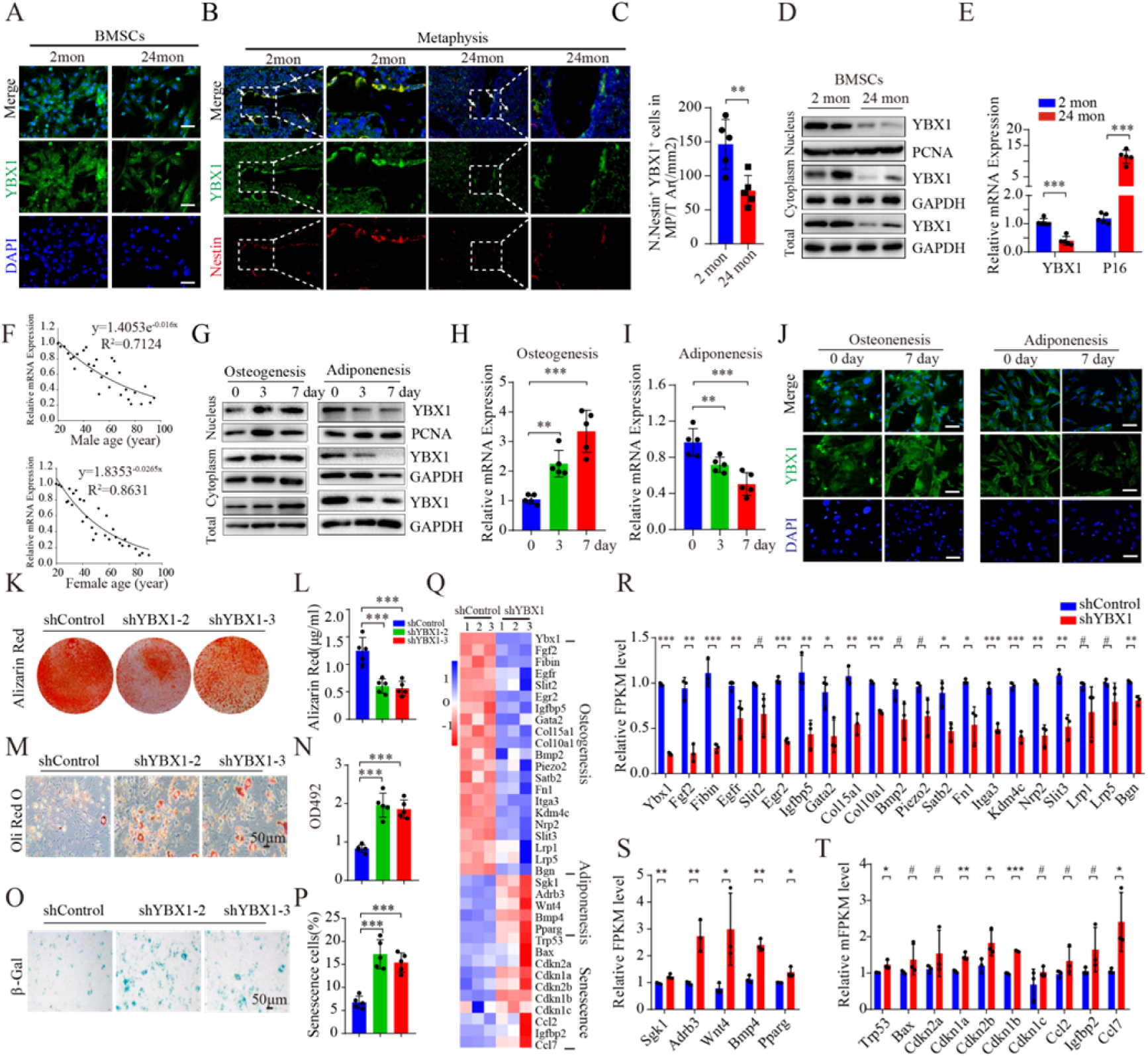
Splicing factor YBX1 regulated fate decision and senescence of BMSCs and showed decreased expression during aging. (A) Representative images of immunofluorescence staining of YBX1 (green) in primary BMSCs. Scale bar: 100μm. (B-C) Representative immunohistochemical staining image of YBX1 (green) and Nestin (red) (B) and quantification of YBX1^+^ Nestin^+^ cells number (C) in femoral bone marrow. Scale bar: 100 μm. (D) Western blot analysis of the expression of YBX1 in BMSCs isolated from mice with different age. (E) qRT-PCR analysis of the expression of *Ybx1* and *P16* in BMSCs isolated from mice with different age. (F) Age-associated changes of YBX1 levels in human BMSCs from 30 males (up panel) and 30 females (down panel). (G) Western blot analysis of the expression of YBX1 in cultured BMSCs during osteogenic differentiation and adipogenic differentiation. (H and I) qRT-PCR analysis of the expression of *Ybx1* in cultured BMSCs during osteogenic differentiation (H) and adipogenic differentiation (I). (J) Representative images of immunofluorescence staining of YBX1 (green) in BMSCs during osteogenic and adipogenic differentiation. Scale bar: 100μm. (K and L) Representative images of Alizarin Red staining (K) and quantification of calcification (L) by detecting the amount of Alizarin Red extracted from the matrix in BMSCs transfected with adenovirus driven control and YBX1 shRNA at 21 days of osteogenic induction. (M and N) Representative images of Oil Red O staining (M) and quantification of lipid formation by detecting the amount of Oil Red O extracted from the matrix (N) in BMSCs at 10 days of adipogenic induction. Scale bar: 50 μm. (O and P) Representative images of β-Gal staining (O) and quantification (P) of β-Gal positive cells in BMSCs. Scale bar: 50 μm. (Q) Heat map of differentially expressed genes between BMSCs transfected with adenovirus driven control and YBX1 shRNA. (R-T) Relative FPKM level of the expression of osteogenic differentiation related genes (R), adipogenic differentiation related genes (S) and senescence related genes (T) between BMSCs transfected with adenovirus driven control and YBX1 shRNA. Data shown as mean ± SEM. ^#^, no significant difference; *, *P* < 0.05; **, *P* < 0.01; ***, *P* < 0.001; Student’s t test for C, E, R-T and One-way ANOVA for H-I, L, N, P.

Reduced osteogenic differentiation tendency and enhanced adipogenic differentiation tendency are the characteristics of aging BMSCs. We found that the RNA and protein level of YBX1 up-regulated during osteogenic induction and down-regulated during adipogenic induction (Figure 2 G-J). To further investigate the role of YBX1 in BMSCs differentiation and senescence, we used adenovirus mediated shRNA to knock down YBX1 in BMSCs and verified the knockdown efficiency by WB tests (Supplemental Figure 1 D). BMSCs with depletion of YBX1 showed lower osteogenic capacity after osteogenic induction indicated by alizarin red staining (Figure 2 K, L) and higher adipogenic differentiation ability after adipogenic induction indicated by Oil Red O staining (Figure 2 M, N). In addition, BMSCs with depletion of YBX1 had higher percentage of β-Gal positive senescence cells compared with the control group (Figure 2 O, P). RNA-Seq analysis also showed that osteogenic differentiation related genes were down-regulated but adipogenic differentiation and senescence related genes were up-regulated in BMSCs with depletion of YBX1 in comparison with the control group (Figure 2 Q-T). Taken together, these results suggested that YBX1 play a role in the regulation of BMSCs senescence and fate decision.

### 3. Depletion of YBX1 in BMSCs resulted in accelerated bone loss and bone marrow fat accumulation

To further investigate the role of YBX1 on the fate decision and senescence of BMSCs in vivo, we crossed *Prx1-cre* transgenic mice with *YBX1^flox/flox^* mice to specifically knock out YBX1 in BMSCs (*YBX1^Prx1-CKO^*). We confirmed the knockout efficiency in BMSCs by qRT-PCR analysis (Supplemental Figure 1 E). RNA-Seq analysis and GO analysis suggested that BMSCs isolated from *YBX1^Prx1-CKO^* mice had altered genes expression which mainly enriched in osteoblast differentiation, bone mineralization, bone/skeletal system development and fat cell differentiation in comparison with the control group (Supplemental Figure 1 F).

Micro-computed tomography (μCT) analysis of the distal femur metaphysis revealed that bone volume was significantly lower in *YBX1^Prx1-CKO^* male mice relative to their *YBX1^flox/flox^* littermates at 3 months and 12 months old (Figure 3 A, B). Additionally, depletion of YBX1 in BMSCs reduced the trabecular thickness and number, while increased the trabecular separation (Figure 3 C-E). Histochemistry and immunohistochemical analysis showed that *YBX1^Prx1-CKO^* mice had significantly higher number of adipocytes in the bone marrow (Figure 3 F, G), but lower number of osteoblasts on the trabecular bone surfaces (Figure 3 H, I). Calcein double labeling revealed that *YBX1^Prx1-CKO^* mice had significant lower trabecular bone mineral apposition rates (MAR) compared with their *YBX1^flox/flox^* littermates (Figure 3 J, K). Constantly, the bone volume, trabecular thickness and number was also significantly lower while the trabecular separation was significantly higher in *YBX1^Prx1-CKO^* female mice relative to their *YBX1^flox/flox^* littermates at 3 months old (Supplementary Figure 2 A-E). These results suggested that depletion of YBX1 accelerating bone loss and stimulating bone marrow fat accumulation.

**Figure 3.**
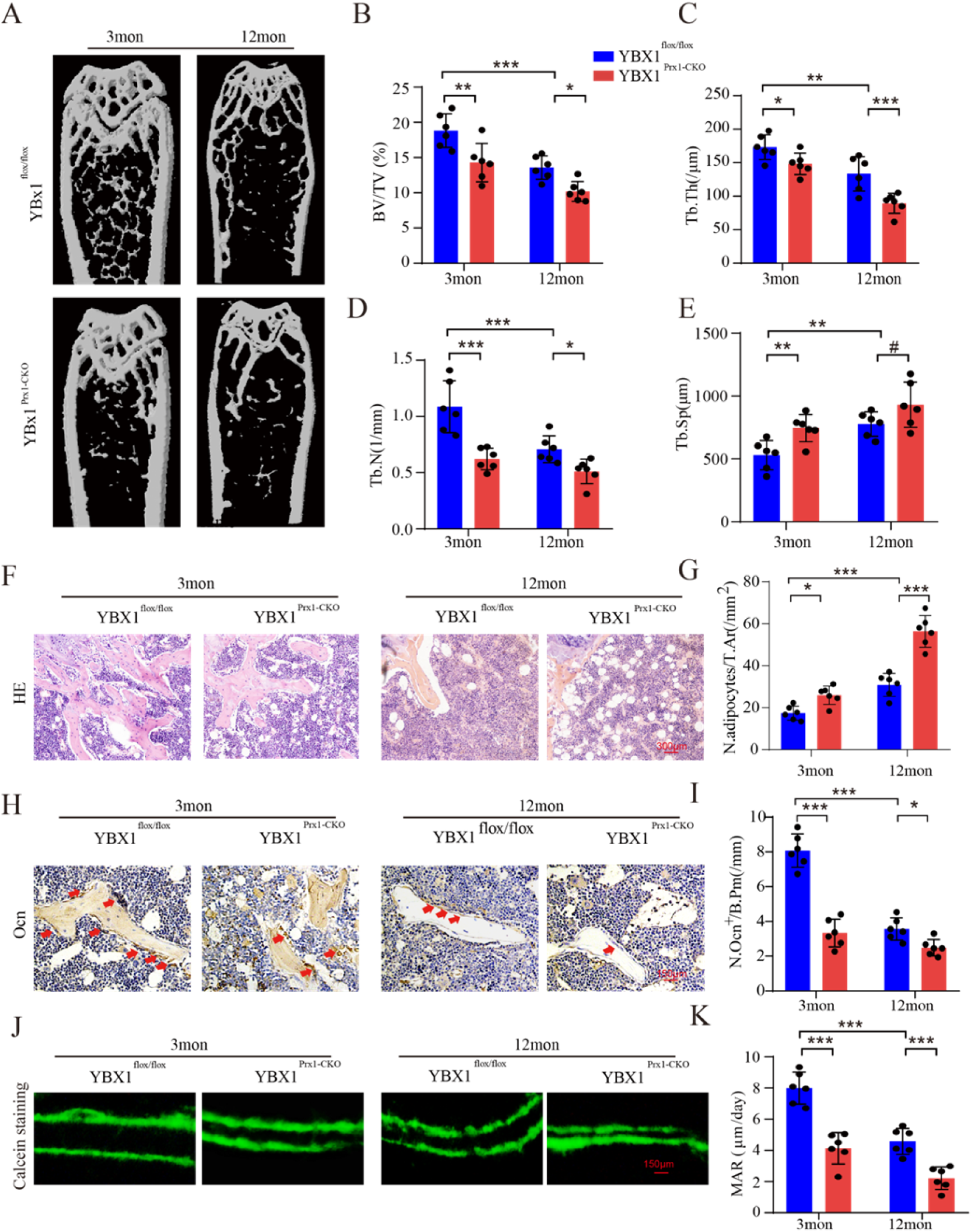
Depletion of YBX1 in BMSCs results in accelerated bone loss and bone marrow fat accumulation. (A-E) Representative μCT images (A) and quantitative μCT analysis of trabecular bone microarchitecture (B-E) in distal femora from 3- and 12-month-old male *YBX1^Prx1-CKO^* mice and *YBX1^flox/flox^* mice. (F) Representative images of HE staining in distal femora. Scale bar: 300μm. (G) Quantification of the number of adipocytes related to tissue area (N. adipocytes/T.Ar). (H) Representative images of osteocalcin (OCN) immunohistochemical staining in distal femora. Arrows point to osteocalcin positive cells. Scale bar: 150μm. (I) Quantification of osteocalcin positive cells in bone surface. Number of OCN^+^ cells per bone perimeter (N. Ocn^+^/B.Pm). (J) Representative images of calcein double labeling of trabecular bone. Scale bar: 150μm. (K) Quantification of mineral apposition rates (MARs). Data shown as mean ± SEM. ^#^, no significant difference; *, *P* < 0.05; **, *P* < 0.01; ***, *P* < 0.001; One-way ANOVA.

### 4. Over-expression of YBX1 attenuated fat accumulation and promoted bone formation in aged mice

Next, we investigated whether elevated the level of YBX1 could attenuated the senescence of BMSCs and stimulated its osteogenic differentiation. BMSCs with over-expression of YBX1 showed enhanced osteogenic differentiation indicated by Alizarin Red staining (Figure 4 A) and restrained adipogenic differentiation indicated by Oil Red O staining (Figure 4 B). Over-expression of YBX1 also reduced BMSCs senescence indicated by β-Gal staining (Figure 4 C). Western blot analysis showed that BMSCs with over-expression of YBX1 had higher level of osteogenic related protein SP7, lower level of adipogenic related protein PPARGγ and lower level of senescence related protein P16 (Figure 4 D).

**Figure 4.**
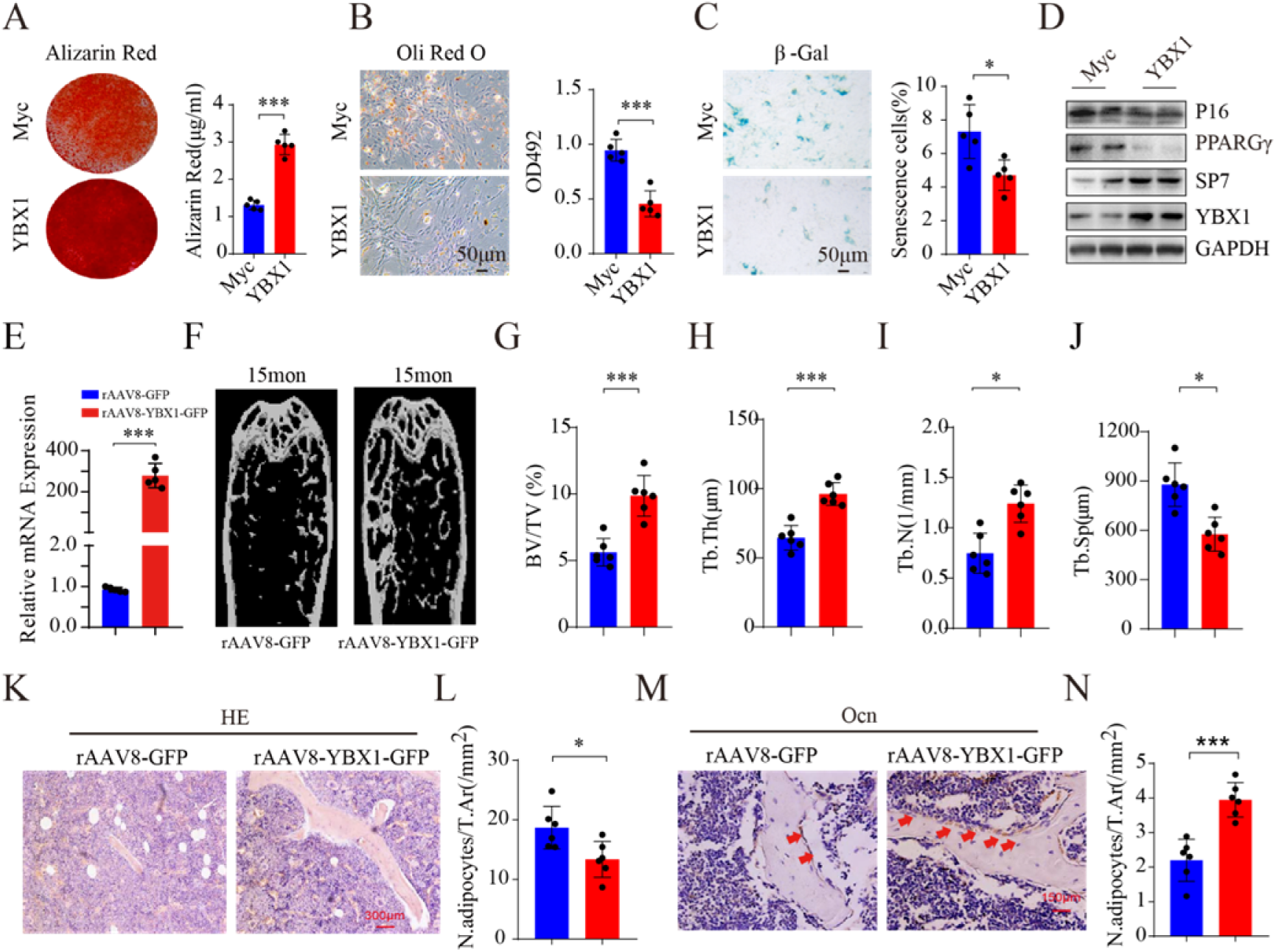
Over-expression of YBX1 attenuated fat accumulation and promoted bone formation in aged mice. (A) Representative images of Alizarin Red staining (left panel) and quantification of calcification (right panel) by detecting the amount of Alizarin Red extracted from the matrix in BMSCs transfected with control or YBX1 plasmid at 21 days of osteogenic induction. (B) Representative images (left panel) and quantification (right panel) of Oil Red O staining in BMSCs transfected with control or YBX1 plasmid at 10 days of adipogenic induction. Scale bar: 50 μm. (C) Representative images of β-Gal staining (left panel) and quantification (right panel) of β-Gal positive cells in BMSCs transfected with control or YBX1 plasmid. Scale bar: 50 μm. (D) Western blot analysis of the expression of YBX1, SP7, PPARGγ and P16 in BMSCs. (E) qRT-PCR analysis of the expression of YBX1 in BMSCs from mice with AAV8-YBX1-GFP or AAV8-GFP transfection. (F-J) Representative μCT images (F) and quantitative μCT analysis of trabecular bone microarchitecture (G–J) in distal femora from 15-month-old mice with AAV8-YBX1-GFP or AAV8-GFP transfection. (K) Representative images of HE staining in distal femora. Scale bar: 300μm. (L) Quantification of the number of adipocytes related to tissue area (N. adipocytes/T.Ar). (M) Representative images of osteocalcin immunohistochemical staining in distal femora. Arrows point to osteocalcin positive cells. Scale bar: 150μm. (N) Quantification of osteocalcin positive cells in bone surface. Number of OCN^+^ cells per bone perimeter (N. Ocn^+^/B.Pm). Data shown as mean ± SEM. ^#^, no significant difference; *, *P* < 0.05; **, *P* < 0.01; ***, *P* < 0.001; Student’s t test.

To investigate whether recovering of YBX1 expression could further alleviate aged-associated bone loss in vivo, we constructed AAV serotype 8 with CMV promoter for gene delivery of YBX1 (rAAV8-YBX1-GFP) to over-express YBX1 in BMCSs. We treated 14-month-old mice with rAAV8-YBX1-GFP by intra–bone marrow injection. One months after, YBX1 expression level in BMSCs isolated from mice infected with rAAV8-YBX1-GFP was much higher than that from control mice (Figure 4 E), μCT analysis showed higher bone volume, trabecular number and lower trabecular separation in YBX1 over-expression mice compared to the control group (Figure 4 F-J). rAAV8-YBX1-GFP treated mice had significant lower bone marrow fat accumulation (Figure 4 K, L) and more osteoblasts on the trabecular bone surfaces (Figure 4 M, N). These results suggested that recovering of YBX1 expression restrained age-related bone loss and bone marrow fat accumulation in old mice.

### 5. YBX1 regulated the fate of BMSCs through regulating the splicing of pre-mRNAs critical for differentiation and senescence

To further investigate the role of splicing factor YBX1 in BMSCs, we evaluated changes in pre-mRNA splicing between BMSCs isolated from *YBX1^Prx1-CKO^* mice and their *YBX1^flox/flox^* littermate controls by calculating “percentage spliced in” (ΔPSI) values of major alternative splicing events (Supplementary Figure 3 A, B). There were 234 pre-mRNAs showed altered splicing with ΔPSI>0.1 and P value<0.05. The change in pre-mRNA splicing upon YBX1 deletion was alternative first exon (45.53%), alternative 5’ splice site (17.45%), exon skipping (13.62%), intron retention (10.21%), alternative 3’ splice site (9.79%) and alternative last exon (3.4%) (Supplementary Figure 3 B). We next performed mass spectrometry (MS) following immunoprecipitation in BMSCs to identify YBX1 interaction partners (Supplementary Figure 3 C). Proteomic network analysis showed that YBX1 interacted with a number of proteins related to ribosome, ribosome biogenesis and spliceosome complex (Supplementary Figure 3 D, E). Protein correlation analysis indicated that YBX1 and its related protein form a regulatory network whose functions are mainly enriched in various RNA metabolic processes including spliceosome and ribonucleoprotein complex assembly, snRNA processing, alternative mRNA splicing and RNA splicing (Supplementary Figure 3 E, F). Among these YBX1 interaction partners, a cluster of ribonucleoproteins, mRNA splicing factors and ribosomal proteins were significantly enriched to take part in spliceosome assembly reaction to form a mature mRNA (Supplementary Figure 3 G, H), which also indicated the core role of splicing factor YBX1 in pre-mRNA altered splicing event.

To further investigate the mechanism by which splicing factor YBX1 regulating the differentiation and senescence of BMSCs, we identified genome-wide targets of YBX1 in BMSCs by anti-YBX1 ultraviolet crosslinking immunoprecipitation (CLIP) analysis using BMSCs cell line (Figure 5 A). CLIP analysis identified 7890 YBX1-biding sites and approximately 51.69% of them were distributed in exons (Figure 5 B), with obvious preferential occupancy of CLIP sequence (Figure 5 C). By combine RNA sequencing and anti-YBX1 CLIP analysis, we identified 66 pre-mRNAs in BMSCs with YBX1-biding sites showed alternative splicing upon YBX1 deletion (Figure 5 D). Among those mRNAs, BMSCs osteogenesis related genes *Fn1* and *Sp7*, BMSCs senescence related gene *Sirt2*, and BMSCs differentiation transcriptional modulate gene *Taz* ^37^, were identified as direct YBX1-mRNA binding targets and went through mis-splicing, including alternative first exon of Fn1, exon skipping of *Sirt2, Sp7* and *Taz* upon YBX1 deletion (Figure 5 E-H). We constructed an RNA map for YBX1-dependent splicing regulation and found that repression and activation related binding occurred at almost completely different sites (Figure 5 I). Semi-quantitative PCR validated the exon skipping of *Sirt2, Sp7* and *Taz* in BMSCs isolated from 24-month-old mice and *YBX1^Prx1-CKO^* mice (Figure 5 J-M). To further investigated whether those mis-splicing would affect BMSCs differentiation, we transfected the different mRNA isoforms of *Taz*, *Sp7* or *Sirt2* into BMSCs which underwent osteogenic or adipogenic induction. The long isoform of *Sp7* mRNA (without skipping of e2) had better promoting effect on osteogenic differentiation in BMSCs (Figure 5 O), the long isoform of *Taz* mRNA (without skipping of e9) had better promoting effect on osteogenic differentiation and better suppression effect on adipogenic differentiation in BMSCs (Figure 5 N, P) and the long isoform of *Sirt2* mRNA (without skipping of e2) had better suppression effect on senescence of BMSCs (Figure 5 Q).

**Figure 5.**
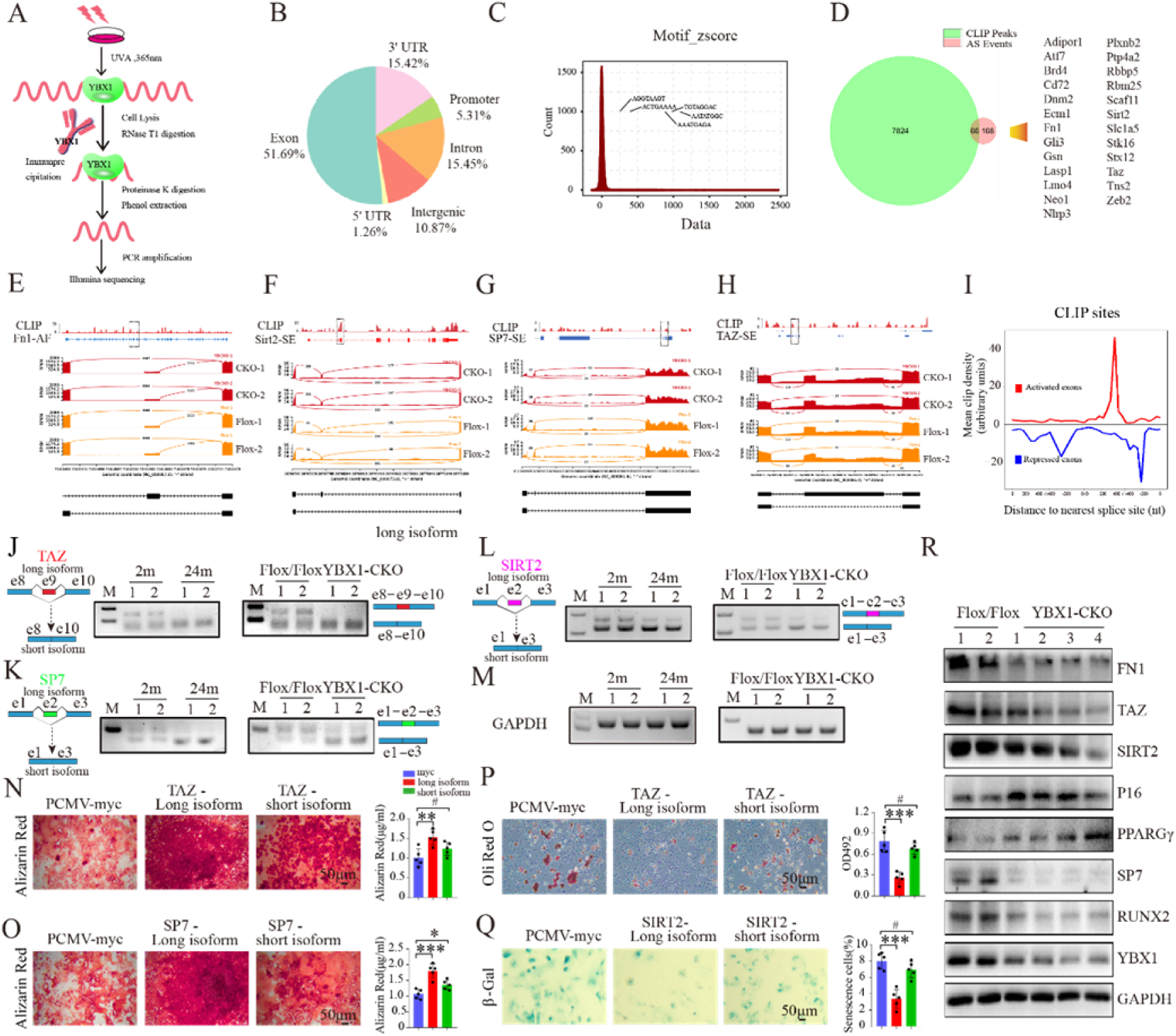
YBX1 regulated the fate of BMSCs through regulating the splicing of pre-mRNAs critical for differentiation and senescence. (A) Schematic diagram of experimental process of anti-YBX1 CLIP analysis. (B) Genomic distribution of YBX1 CLIP-seq peaks. (C) Enriched motifs for YBX1 binding. Inset shows consensus sequence, deduced from the top ten motifs. (D) Venn diagrams of overlapping genes targeted by YBX1 and showed alternative splicing upon YBX1 deletion. (E-H) RNA-seq read coverage across *Fn1* (E), *Sirt2* (F), *Sp7* (G) and *Taz* (H) from BMSCs isolated from *YBX1^Prx1-CKO^* mice and *YBX1^flox/flox^* mice. (I) Mean CLIP density near exons repressed (blue), activated (red) by YBX1. (J-M) Semi-quantitative PCR showed the different isoforms of *Taz* (J)*, Sp7* (K), *Sirt2* (L) and reference genes *Gapdh* (M) between BMSCs isolated from 2-month-old or 24-month-old mice and from *YBX1^Prx1-CKO^* mice or *YBX1^flox/flox^* mice. (N-O) Representative images of Alizarin Red staining (left panel) and Quantification of calcification by detecting the amount of Alizarin Red extracted from the matrix (right panel) in BMSCs transfected with different isoforms of *Taz* (N) or different isoforms of *Sp7* (O). (P) Representative images (left panel) and quantification of Oil Red O staining (right panel) in BMSCs transfected with different isoforms of *Taz* with 10 days of adipogenic induction. (Q) Representative images of β-Gal staining (left panel) and quantification (right panel) of β-Gal positive cells in BMSCs transfected with different isoforms of *Sirt2.* (R) Western blot analysis of the expression of YBX1, FN1, TAZ, SIRT2, P16, PPARGγ, SP7 and RUNX2 in BMSCs isolated from *YBX1^Prx1-CKO^* mice and *YBX1^flox/flox^* mice. Data shown as mean ± SEM. ^#^, no significant difference; *, *P* < 0.05; **, *P* < 0.01; ***, *P* < 0.001; One-way ANOVA.

In order to evaluate whether those altered splicing events in pre-mRNA would result in a variation in protein level, we performed WB test and demonstrated a decreased expression in FN1, TAZ, SIRT2 and SP7, which pre-mRNA had direct YBX1-mRNA binding target and went through mis-splicing, in BMSCs isolated from *YBX1^Prx1-CKO^* mice (Figure 5 R). TAZ was reported to form a transcriptional complex with RUNX2 that drives osteogenic differentiation of BMSCs, coordinately represses adipocyte differentiation in a transcriptional repressor of PPARGγ^14,37^. Our previous research suggested that YBX1 could directly bind to the promoter and repress the expression of P16 in hypothalamic neural stem cells^27^. We also detected a decreased expression in RUNX2 and increased expression level of P16 and PPARGγ in BMSCs isolated from *YBX1^Prx1-CKO^* mice (Figure 5 R).

In nucleus, YBX1 mainly performs its pre-mRNA splicing function, meanwhile YBX1 also could bind to the 3’UTR region of mRNA to maintain its stability in cytoplasm ^26^. We also detected many binding sites of YBX1 in 3’UTR region of mRNA in BMSCs (Figure 5 B). The protein level of YBX1 was also lower in cytoplasm of BMSCs isolated from older mice and in cultured primary BMSCs from later passage (Figure 2 D and Supplemental Figure 1 B). By combine RNA sequencing and anti-YBX1 CLIP analysis, we identified 89 mRNAs in BMSCs with YBX1-biding sites on 3’ UTR region showed altered expression upon YBX1 deletion (Supplemental Figure 4 A, B). Among those mRNAs, senescence related gene *Nrp2*, osteogenesis related genes including *Bgn*, *Colla2* and *Thbs1* with its downstream FAK signaling, showed decreased expression upon YBX1 deletion (Supplemental Figure 4 C-E),

As an RNA binding protein, YBX1 also could bind to the promoter regions of genes and regulated their expression ^28,38^. To investigate whether YBX1 regulated those senescence and osteogenesis genes of BMSCs at transcriptional level, we performed YBX1 chromatin immunoprecipitation sequencing (ChIP-seq) and found that only 2.69%peaks located at the promoter region (Supplemental Figure 5 A) and there was no significant difference between the enrichment of YBX1 and input at the transcription start sites TSSs (Supplemental Figure 5 B). Additionally, there was no significant binding of YBX1 to promoter region of *Bgn, Colla2, Nrp2, Thbs1 Fn1, Taz, Sirt2*, and *Sp7* genes (Supplemental Figure 5 C, D).

Taken together, these results suggested that YBX1 regulated the expression level of osteogenic differentiation, adipogenic differentiation and senescence related genes in BMSCs by controlling pre-mRNA alternative splicing and mRNA stability.

### 6. Sciadopitysin bind to and inhibited ubiquitin degradation of YBX1

To search the potential therapeutic strategy to restrain the age-related debility of BMSCs by targeting YBX1, we performed molecular docking to screen the natural small molecular compounds that interacted with mouse YBX1 as previously reported ^27^. We chose 9 top-ranked small molecules which are related to anti-oxidation, anti-inflammation, anti-aging and bone metabolism (Figure 6 A). Among these candidates, 5 compounds including theaflavin-3-gallate, eriocitrin, sciadopitysin, isoginkgetin and bilobetin showed no adverse effect on BMSCs proliferation evaluated by CCK8 assay (Figure 6 B) and all of these 5 compounds had no effects on the transcription of *YBX1* (Figure 6 C). Only sciadopitysin and theaflavin-3-gallate could promote osteogenic differentiation, inhibit adipogenic differentiation and attenuate senescence of BMSCs, between them, sciadopitysin showed better effects (Figure 6 D-G). So, we choose sciadopitysin for further study. The structure and binding mode of sciadopitysin and YBX1 showed that sciadopitysin could enter into the pocket-like structure of YBX1 (Figure 6 H). Different dose of sciadopitysin increased the protein level of YBX1 in BMSCs in a graded manner (Figure 6 I). Sciadopitysin treatment could also increase the expression of FN1, TAZ, SP7, THBS1 in BMSCs at the protein level (Figure 6 I).

**Figure 6.**
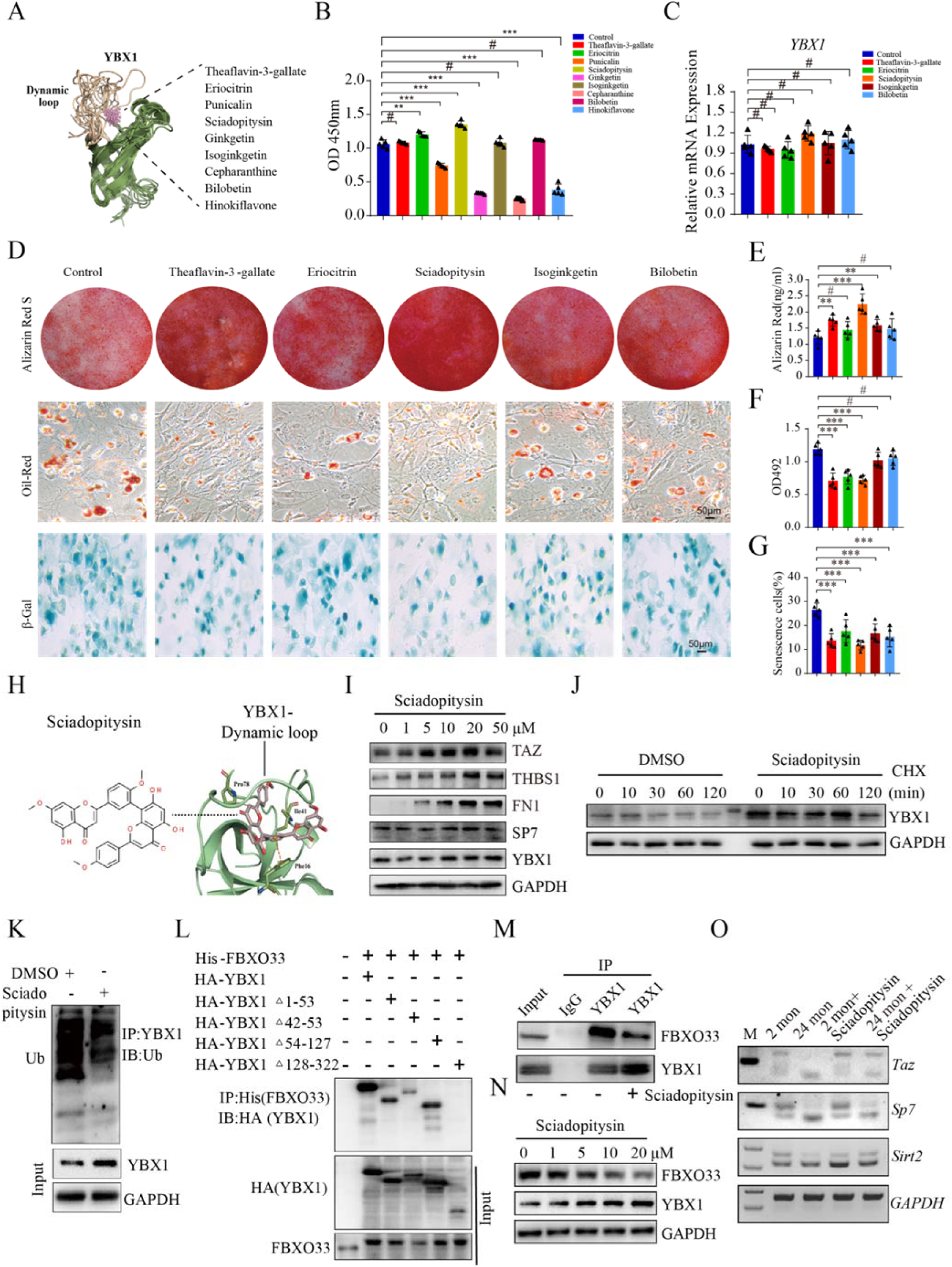
Sciadopitysin bind to and inhibited ubiquitin degradation of YBX1. (A) The homology modeling structure of mouse YBX1 and 9 top-ranked candidates. (B) Cell proliferation rate was assessed by CCK8 assay with administration of different compounds. (C) qRT-PCR analysis of *Ybx1* expression in BMSCs with administration of different compounds. (D) Representative images of Alizarin Red staining (up panel), Oil Red O staining (middle panel, scale bar: 50 μm.) and β-Gal staining (bottom panel, scale bar: 50 μm.) in BMSCs treated with different compounds. (E-G) Quantification of calcium mineralization based on Alizarin Red staining (E), quantification of Oil Red O based on Oil Red O staining (F) and quantification of β-Gal positive cells based on β-Gal staining in BMSCs treated with different compounds (G). (H) The molecular structure of sciadopitysin and model of interaction between sciadopitysin and mouse YBX1. (I) Western blot analysis of TAZ, THBS1, FN1, SP7, YBX1 expression in BMSCs treated with different concentration of sciadopitysin. (J) Western blot analysis of YBX1 in sciadopitysin pre-treated BMSCs with treatment of cycloheximide CHX. (K) Western blot analysis of YBX1 related ubiquitination in sciadopitysin pre-treated BMSCs with Mg132. (L) Co-IP of His-FBXO33 with HA-YBX1 and serious of mutant HA-YBX1 following transfection into BMSCs. (M) Co-IP of FBXO33 withYBX1 with or without administration of sciadopitysin. (N) Western blot analysis of FBXO33 and YBX1 expression in BMSCs treated with different concentration of sciadopitysin. (O) Semi-quantitative PCR showed the isoforms of *Sirt2, Sp7* and *Taz* in cultured BMSCs isolated from 2-month-old or 24-month-old mice then treated with or without sciadopitysin. Data shown as mean ± SEM. ^#^, no significant difference; **, *P* < 0.01; ***, *P* < 0.001; One-way ANOVA.

To investigate the means by which sciadopitysin treatment increased the protein level of YBX1, we blocked the protein synthesis in BMSCs by cycloheximide (CHX) and found that sciadopitysin treatment slow down the degradation of YBX1 (Figure 6 J). YBX1 was demonstrated to be degraded through ubiquitination ^39^. Therefore, we further tested whether sciadopitysin treatment affected YBX1 ubiquitination and found that sciadopitysin significant decreased the ubiquitination level of YBX1 (Figure 6 K). Previous studies reported that ubiquitin ligase FBXO33 could bind to and mediate the ubiquitination of YBX1^39^. To confirmed the interaction between FBXO33 and YBX1 and investigate which region of YBX1 protein could bind to FBXO33, we generated a series of YBX1 plasmid mutants, transfected them into BMSCs and performed co-IP assay. The results showed that deletion of the amino acid cites 128-322 (C-terminus) and the amino acid cites 42-53 (pocket area) of YBX1 impaired the interaction between YBX1 and FBXO33, suggesting that the C-terminus and the pocket area is crucial for YBX1 binding to FBXO33 (Figure 6 L). We further confirmed the interaction between FBXO33 and YBX1 in BMSCs by IP test, and the interaction was suppressed by sciadopitysin treatment (Figure 6 M). Notably, sciadopitysin treatment could decreased the expression level of FBXO33 in BMSCs (Figure 6 N). These results suggested that sciadopitysin inhibit ubiquitination degradation of YBX1 both by decreased the level of ubiquitin ligase FBXO33 and prevent YBX1 from combining with FBXO33. Furthermore, sciadopitysin treatment also could partially restrain the exon skipping of *Sirt2, Sp7* and *Taz* in BMSCs isolated from 24-month-old mice (Figure 6 O), which indicated that sciadopitysin could alleviate age related mis-splicing of destiny genes in aged BMSCs.

### 7. Sciadopitysin treatment ameliorated age-related bone loss in mice

To further investigate whether sciadopitysin administration could alleviate age-related bone loss, 13-month-old C57/BL6J mice were administrated with sciadopitysin at 40mg/kg body weight per day or with vehicle for 2 months (Figure 7 A). μCT analysis showed that sciadopitysin treated mice had higher bone volume, trabecular thickness, trabecular number and lower trabecular separation compared to vehicle treated and control mice (Figure 7 B-C). Sciadopitysin treated mice had significantly higher number of osteoblasts on the trabecular bone surfaces, as compared with control and vehicle-treated mice (Figure 7 D). Moreover, calcein double labeling analysis showed that sciadopitysin treated mice had significantly higher trabecular bone mineral apposition rates (MAR) compared with control and vehicle-treated mice (Figure 7 E). The number of adipocytes were also decreased by the treatment of sciadopitysin (Figure 7 G). However, the number of osteoclasts and adipocytes were not affected by the treatment of sciadopitysin (Figure 7 F). These results suggested that mice treated with sciadopitysin increased bone formation in old mice.

**Figure 7.**
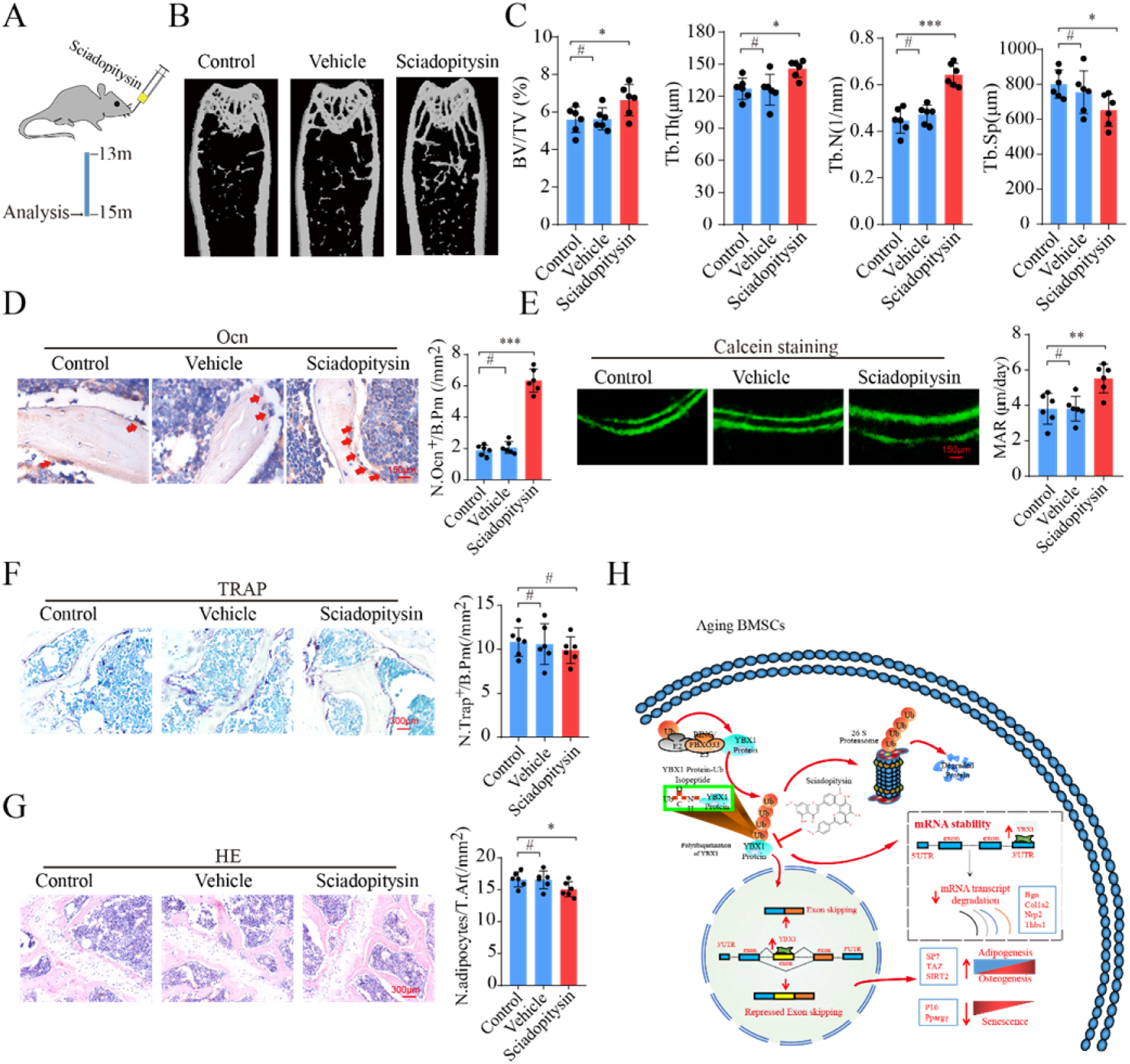
Sciadopitysin treatment alleviates aging-related bone loss in mice. (A) Schematic of the time of oral treatment of sciadopitysin in mice. (B and C) Representative μCT images (B) and quantitative analysis of trabecular bone microarchitecture (C) in distal femora of 15-month-old mice with administration of sciadopitysin or vehicle. (D) Representative images (left panel) and quantification (right panel) of osteocalcin positive cells in distal femora of 15-month-old mice with administration of sciadopitysin or vehicle. Number of Ocn^+^ cells per bone perimeter (N. Ocn^+^/B.Pm). Arrows point to osteocalcin positive cells. Scale bar: 150μm. (E) Representative images (left panel) of calcein double labeling of trabecular bone and quantification (right panel) of mineral apposition rates (MARs) of 15-month-old mice with administration of sciadopitysin or vehicle. Scale bar: 150μm. (F) Representative images (left panel) and quantification (right panel) of Trap positive cells in distal femora of 15-month-old mice with administration of sciadopitysin or vehicle. Number of Tarp^+^ cells per bone perimeter (N. Trap^+^/B.Pm). Scale bar: 300μm. (G) Representative images of HE staining in distal femora (left panel) and quantification of the number of adipocytes related to tissue area (right panel, N. adipocytes/T.Ar) in distal femora of 15-month-old mice with administration of sciadopitysin or vehicle. Scale bar: 300μm. (H) RNA binding protein YBX1 regulate cluster of genes including *Fn1, Taz, Sirt2, Sp7* as a splicing factor in nucleus and *Bgn, Colla2, Nrp2* and *Thbs1* as mRNA stabilized protein in cytoplasm, which further stimulate osteogenic differentiation and restrain senescence of BMSCs. The decreased expression level of YBX1 during aging contribute to the debility of BMSCs including increased senescence and reduced osteogenesis. Sciadopitysin can delay the ubiquitination degradation of YBX1 by preventing YBX1 from binding to ubiquitin ligase FBXO33. (Model based on data from previous figures.) Data shown as mean ± SEM. ^#^, no significant difference; *, *P* < 0.05; **, *P* < 0.01; ***, *P* < 0.001; One-way ANOVA.

Taken together, we demonstrated that RNA binding protein YBX1 can stimulate osteogenic differentiation and restrain senescence of BMSCs by regulating a cluster of genes including *Fn1, Taz, Sirt2, Sp7* as a splicing factor in nucleus and regulating *Bgn, Colla2, Nrp2 and Thbs1* as mRNA stabilized protein in cytoplasm. The decreased expression level of YBX1 during aging contribute to the debility of BMSCs including increased senescence and reduced osteogenesis. Moreover, we identified a natural small compound, sciadopitysin, which can delay the degradation of YBX1 and attenuate age-related bone loss (Figure 7H).

## Discussion

Age-related dysfunction of BMSCs such as lineage switching between osteogenic and adipogenic fates and acceleration in senescence are critical in age-associated osteoporosis. Alternative splicing, as an important regulation pathway of gene translation, has a wide range of biological functions. Disruption in alternative splicing can lead to dysfunction or disease^40^. Aberrant of alternative splicing can accelerate cellular senescence^41^, disrupt the differentiation of mesenchymal stem cells^18–20^. It has been reported that MSC from old and young donors have different alternative splicing events^42^. However, whether aberrant of alternative splicing take part in the aging related dysfunction of BMSCs and how does it work remains unclear. In the present study we observed altered splicing events and changed gene expression pattern of BMSCs in mice during aging by whole transcriptome resequencing analysis and alternative splicing analysis. We further demonstrated that the deficiency of splicing factor YBX1 might be responsible for mis-splicing on BMSCs destiny genes and further result in senescence and change of differentiation direction in aged BMSCs.

As well-known transcriptional and translational regulator, YBX1 take part in variety of RNA-dependent events, including pre-mRNA transcription and splicing, mRNA packaging, mRNA stability and translation ^23,43^. At the cell level, the activities of YBX1 involve in the regulation of multiple processes of cellular biology, such as cell proliferation, differentiation, stress response, and malignant cell transformation^29,33,38,43^. In this study, we found the expression level of YBX1 in BMSCs was decreased during aging, and further demonstrated that BMSCs osteogenesis related genes *Fn1* and *Sp7*, senescence related gene *Sirt2*, and *Taz* which has been reported both involved in osteogenic and adipogenic differentiation of BMSCs ^14,37^, were identified as direct YBX1-mRNA binding targets and went through altered splicing. As a multifunctional RNA binding protein, YBX1 not only act as a splicing factor but also through other ways to regulate BMSCs fate decisions. Our previous research suggested that YBX1 could directly bind to the promoter and repress the expression of *P16*, and thus controlling the senescence of hypothalamic neural stem cells ^28^. In this study we also identified 89 mRNAs in BMSCs, including several osteogenesis related genes and senescence related genes with YBX1-biding sites on 3’ UTR region which showed altered expression upon YBX1 deletion. These results suggested that beside pre-mRNA alternative splicing, YBX1 also could regulate BMSCs fate decisions by directly regulating the transcription of specific genes or maintaining the stability of mRNAs. YBX1 involving the regulation of BMSCs fate decide during aging, and pre-mRNA alternative splicing is one of the important regulatory approaches.

Through further research, we found that BMSC specific YBX1 knockout mice had accelerated bone loss and bone marrow fat accumulation than the control mice, over-expression of YBX1 in bone marrow with AAV8-CMV-YBX1-GFP attenuated bone loss and bone marrow fat accumulation, which further confirmed that YBX1 is a critical factor in orchestrating lineage commitment of BMSCs during aging. Our finding indicated that restore the level of YBX1 in BMSCs during aging might be a therapeutic strategy to alleviate age-associated osteoporosis.

Sciadopitysin is an amentoflanove-type biflavonoid, which is contained in taxus chinensis. Eun Mi Choi et al. using MC3T3-E1 cell lines demonstrated that sciadopitysin protect osteoblast function via upregulation of mitochondrial biogenesis. ^44,45^. In our study, we identified sciadopitysin that could bind to YBX1 to attenuate its ubiquitination degradation, and further increase osteogenic differentiation and inhibit senescence of BMSCs. Treatment of sciadopitysin could attenuate age-related bone loss in mice and we also observed that sciadopitysin treatment could partially restrain the exon skipping of *Sirt2, Sp7* and *Taz* in BMSCs isolated from aged mice. All these results indicated that sciadopitysin perform its protection effect on bone mass partly through YBX1. In this study we didn’t investigate other mechanisms involved and couldn’t exclude the effect of sciadopitysin outside of YBX1, and the efficacy and safety of sciadopitysin need to be further evaluated in larger animals. We believe that sciadopitysin could act as a major candidate compound when designing of new anti-osteoporosis drugs.

Taken together, our study demonstrated that YBX1 worked as an alternative splicing factor in regulating BMSCs senescence and fate decisions and could be a potential therapeutic target for the treatment of age-related osteoporosis.

## Materials and Methods

### Animals

*YBX1^flox/+^* mice were purchased from Cyagen Biosciences. *Prx1-cre* transgenic mice were purchased from the Jackson Laboratory. We mated *YBX1^flox/+^* male mice with *YBX1^flox/+^* female mice to obtain *YBX1^flox/flox^* mice. We crossed prx1-cre mice with *YBX1^flox/flox^* to obtain *prx1-cre;YBX1^flox/+^* mice. By mating *prx1-cre;YBX1^flox/^*^+^ male mice with *YBX1^flox/flox^* female mice, we obtained *prx1-cre;YBX1^flox/flox^* mice as homozygous conditional YBX1 knockout mice. The littermate *YBX1^flox/flox^* mice were used as controls.

All the mice used in this study were bred under specific-pathogen-free conditions of Laboratory Animal Research Center at Central South University. All the mice were kept in a C57BL/6 background. All animal care protocols and experiments were approved by the Medical Ethics Committee of Xiangya Hospital of Central South University. Approval number: 2019030350.

### Intra-bone marrow injection of adeno-associated virus

Intra-bone marrow delivery of virus was performed as previous reported ^14^. Recombinant adeno-associated serotype 8 virus with CMV promoter for YBX1 overexpression (rAAV8-YBX1-GFP) was purchased from Hanbio Biotechnology Co (Shanghai, China). The virus was diluted with sterile PBS and the viral titer used in the study was 5×10^12^ vg/ml. We used rAAV8-GFP as control. 5 μl of either rAAV8-YBX1-GFP or rAAV8-GFP was delivered into the femoral medullary cavity through periosteal injection twice a month for 2 months.

### Compound treatment

Sciadopitysin (T5S2129), Eriocitrin (T6S0221), Isoginkgetin (T4S21320), Bilobetin (T4S2128), Theaflavin3-gallate (T3051), Punicalin (T4S1718), Ginkgetin (T4S2126), Cepharanthine (T0131) and Hinokiflavone (T4S0181) were purchased from TargetMol. For in vivo studies, sciadopitysin was treated by oral gavage at 40mg/kg body weight/day for 2 months. For in vitro experiment, sciadopitysin, eriocitrin, isoginkgetin, bilobetin, theaflavin3-gallate, punicalin, ginkgetin, cepharanthine and hinokiflavone were dissolved in DMSO and treated at the concentration of 10uM unless specified otherwise.

### Primary mouse BMSC isolation and culture

Primary mouse BMSCs were isolated as previously reported ^15^. We flushed bone marrow cells from the tibia and femora of male mice and incubated the cells with anti-Sca-1-PE (108108; BioLegend), anti-CD29-FITC (102206; BioLegend), anti-CD45-PerCP (103132; BioLegend), and CD11b-PerCP (101226; BioLegend) for 20 minutes at 4°C. For human BMSCs, human bone marrow cells were collected and incubated with FITC-, APC-, and PE-conjugated antibodies that recognized human Stro-1 (BioLegend, 340106), CD45 (BioLegend, 304012), and CD146 (BioLegend, 361008) at 4°C for 30 minutes. The acquisition was conducted on a fluorescence-activated cell sorting (FACS) Aria model (BD Biosciences). FACS DIVE software version 6.1.3 (BD Biosciences) was used for the analysis.

The sorted mouse Sca-1^+^CD29^+^CD45^-^CD11b^-^ BMSCs and human CD146+Stro-1+CD45− BMSCs were cultured for about 1 week to reach 80%–85% confluence. Then, first-passage BMSCs were digested with trypsin for about 1min and seeded in culture dishes for the enrichment of cell populations.

### Cell culture, transfection, differentiation and senescence assay

BMSCs were cultured in α-MEM supplemented with 10% fetal bovine serum (Gibco), 100 μg/ml streptomycin (Gibco) and 100 units/ml penicillin (Gibco) at 37 °C with a humidified atmosphere of 5% CO_2_.

For YBX1 knock down, adenovirus particles of shYBX1 and shControl were obtained from Hanbio Biotechnology Co (Shanghai, China). BMSCs were infected with adenovirus for 8 hours before proceeding to perform further experiments. For YBX1 over-expression, mYBX1 pcDNA3.1 was purchased from Youbio Biological Technology. The mYBX1 plasmid and negative control were transfected into BMCSs using Lipofectamine 2000 (Invitrogen) for 6 hours before proceeding to perform further experiments.

For osteogenic differentiation assay, BMSCs were cultured in 6-well plates at a density of 1.0×10^6^ cells per well with osteogenic induction medium (10 mM β-glycerol phosphate, 0.1 uM dexamethasone, and 50 uM ascorbate-2-phosphate) for 3 weeks. Then, we stained the cells with 2% Alizarin Red (Cyagen Biosciences) to detect the cell matrix calcification. Alizarin Red was extracted from the matrix with cetyl-pyridinium chloride solution and quantified using spectrophotometry at 562 nm. For adipogenic differentiation assay, BMSCs were plated in 6-well plates at a density of 2.5×10^6^ cells per well with adipogenic induction medium (1 μM dexamethasone, 5 μg/ml insulin and 0.5 mM 3-isobutyl-1-methylxanthine) for 10 days. Culture medium was changed every 3 days. Lipid droplets in mature adipocytes were detected by Oil Red O staining according to the manufacturer’s instruction (Cyagen Biosciences). Oil Red O was extracted from the matrix and quantified using spectrophotometry at 492 nm.

For cellular senescence assay, BMSCs were seeded in 6-well plates at a density of 1.0 × 10^6^ cells per well for 24 hours. Senescent cells were stained using a β-Gal staining kit (Solarbio Science Technology) according to the manufacture’ instructions.

### Histochemistry analysis

Histochemistry analysis was performed as previously described ^14,15^. Briefly, After the mice were euthanized, bones were harvested, fixed in 4% paraformaldehyde (PFA) for 24 hours at 4°C, decalcified in 10% EDTA for 3 weeks at 4 °C, embedded in paraffin. 4-μm-thick longitudinal bone sections were made and stained with TRAP (Sigma-Aldrich) and HE (Servicebio) according to the manufacturer’s instructions.

### Immunohistochemical staining

Immunohistochemical staining was performed as previously reported ^46^. Briefly, after antigen retrieval, bone sections were blocked in 5% bovine serum albumin (BSA) for 1 hour at room temperature and incubated with primary antibody to osteocalcin (Takara, M173) at 4°C overnight. Then the bone sections were incubated with appropriate secondary antibody at room temperature for 1 hour. Finally, we detected the immunoactivity with an HRP-streptavidin detection system (Dako), and counterstained the slides with hematoxylin.

### Calcein double-labeling assay

To evaluate dynamic bone formation ability, mice were administrated intraperitoneally with calcein (25 mg/kg, SigmaAldrich) at 8 and 2 days before euthanasia. After fixation in 70% ethanol, the samples were dehydrated in gradient ethanol. Then the calcein double labeled bones were embedded in methyl methacrylate. 5-μm-thick longitudinal bone sections were made using a microtome and observed under a fluorescent microscope.

### Immunofluorescence staining

Cultured BMSCs were fixed with 4% PFA for 15 minutes at room temperature. Then the cells were blocked with 5%BSA for 1 hour at room temperature, and incubated with YBX1 antibody (Cell Signaling Technology, 4202, 1:100), nestin antibody (Millipore, MAB353, 1:100) over night at 4°C. After that, the cells were incubated with Alexa 488 (Invitrogen, A21106) and Alexa 555 (Invitrogen, A21422) conjugated secondary antibodies and the nucleus were stained with Dapi.

For bone sections, after antigen retrieval, bone sections were blocked with 5% BSA for 1 hour at room temperature. Then the bone sections were incubated with YBX1 antibody (Cell Signaling Technology, 4202, 1:100), nestin antibody (Millipore, MAB353, 1:100) over night at 4°C. Next, the bone sections were incubated with Alexa 488 (Invitrogen, A21106) and Alexa 555 (Invitrogen, A21422) conjugated secondary antibodies and the nuclear were stained with Dapi.

### RNA sequencing and analysis

Total RNAs were extracted from shYBX1 and shControl infected BMSCs using Trizol reagent. The NanoPhotometer^®^ spectrophotometer (IMPLEN, CA, USA) was used to check RNA purity. RNA integrity was evaluated by the RNA Nano 6000 Assay Kit of the Bioanalyzer 2100 system. NEBNext® UltraTM RNA Library Prep Kit for Illumina® (NEB, USA) was used to generate sequencing libraries according to manufacturer’s recommendations. We controled the quality of RNA-seq data by removing low quality reads, reads containing ploy-N, reads containing adapter from raw data. We mapped clean data to the reference genome using Hisat2 v2.0.5. The DESeq2 R package (1.16.1) was used to perform differential expression analysis of two groups. The gene expression is considered to be significantly different if displaying≥1.5 fold change and P value < 0.05. Event level differential splicing was calculated with the EventPointer package25 in R.

### CLIP

BMSCs (1*10^8^ cells) were treated with 4-thiouridine (100 μM) for 16h. After 16 h incubation, cells were washed twice with 10 ml of ice-cold PBS and then were UV irradiated at 150 mj/cm2 on ice. Cells were collected with a clear cell scraper and transferred into a new 15 ml centrifuge tube and then pelleted at 1,000 g for 5min at 4°C. The supernatant was discarded and the cell pellet was re-suspended with 12 mL 1×cell lysis buffer containing 120 μL 100×DTT and 120 μL 100× Protease inhibitor and incubated on ice for 10min. Cell lysates were centrifuged at 14,000 g for 15 minutes at 4°C and transfered supernatant into a new 15 ml centrifuge tube. For CLIP procedure, 8ml supernatant was incubated with 5 μg YBX1 antibody (Cell Signaling Technology, 4202) and 4ml supernatant was incubated with 2 μg IgG antibody overnight at 4°C. The next day, CLIP samples were further incubated with 40 μL ProteinA/G magnetic beads for 3h at 4°C. The magnetic beads were washed twice with 1×IP wash buffer and subsequently resuspended in 60 μL 1×IP wash buffer containing 6 μL RNase T1. The magnetic beads were incubated at 22°C for 60min and at ice for 5min and then washed twice with 1×IP wash buffer. Next, the magnetic beads were resuspended in 100 μL 1×IP wash buffer containing 20 μL DNase I and incubated at 37°C for 15min and at ice for 5min. The beads were Place on the magnetic separator then washed twice with 1×IP wash buffer. Finally, the magnetic beads were resuspended in 1×100μl Proteinase K and incubated at 55°C for 30min. To remove the beads by a magnetic separator, and transfer the supernatant into a new 1.5ml centrifuge tube. An equal volume acidic phenol: chloroform: isoamylalcohol (25:24:1) and equal volume chloroform was used to clear the supernatant. After, the clear supernatant was added with 2 volumes of ethanol and one microliter of glycogen and then precipitated at -20°C for 2h. The supernatant was centrifuge at 14,000 rpm for 30 minutes at 4°C and discarded the supernatant. The precipitate was washed twice with 75% ethanol and re-suspended with 10 μL RNase-free water. The recovered RNA was used to perform the high-throughput sequencing with Illumina NextSeq 500 system under the help of ABLife Inc and Wuhan Igenebook Biotechnology Co.,Ltd.

### Liquid chromatography (LC)–MS/MS measurement

We used Q-Exactive mass spectrometer (Thermo Fisher) connected online to an Easy-nLC 1000(Thermo Fisher) to analyze the LC-MS/MS. Separation of peptides was conducted using Zeba Spin columns (Pierce) in a gradient from 5–65% in buffer B (0.1% formic acid, 84% acetonitrile). The columns temperature was kept at 50 °C in an oven. We analyzed the peptides using a full scan (300-1,600 m/z, R = 60,000 at 200 m/z) with 3e6 ions as the target. Then we performed high energy collisional disassociation for fragmentation of top 20 most rich isotype patterns with a charge ≥2 MS/MS scan. All data were collected with X-caliber software (Thermo Fisher).

### ChIP-seq

BMSCs were crosslinked with 1% formaldehyde for 10 min at room temperature and nuclei were extracted, lysed and sheared on ice. Chromatin was diluted with ChIP buffer, cleared and incubated with 5ug YBX1 antibody (Cell Signaling Technology, 4202) overnight at 4°C. The antibody/chromatin complex was immunoprecipitated with protein G beads. Then the antibody/chromatin complex was extensively washed, eluted and de-crosslinked. After purification, ChIP DNA was used to prepare ChIP-sequencing libraries with SimpleChIP^®^ ChIP-seq DNA Library Prep Kit for Illumina^®^ and subjected to sequencing on an Illumina NextSeq platform under the help of Seqhealth Technology Co., LTD (Wuhan, China).

### qRT-PCR analysis

We used Trizol reagent (Invitrogen) to isolate total RNA from cells based on the standard protocol. We conducted reverse transcription with 1ug total RNA with Evo M-MLV RT Kit with gDNA Clean for qPCR AG11705 (Accurate Biotechnology (Hunan) Co., Ltd). Quantitatification of mRNA was detected by qRT-PCR with SYBR^®^ Green Premix Pro Taq HS qPCR Kit ( Rox Plus ) AG11718 (Accurate Biotechnology(Hunan) Co., Ltd).

### Western blot

Western blot was performed as previously described ^47^. The antibodies used for western blot are: YBX1 (D2A11) (Cell Signaling Technology, 9744, 1:1000), β actin (Origene, TA811000, 1:5000), GAPDH (Origene, TA802519, 1:5000), PCNA (BOSTER Biological Technology, BM0104, 1:5000), PPARGγ (81B8)(Cell Signaling Technology, 2443, 1:1000), P16 (Sigma-Aldrich, SAB5300498, 1:1000), Fibronectin (Santa Cruz, sc-8422, 1:500), TAZ (E8E9G) (Cell Signaling Technology, 83669, 1:1000), Sirt2 (Abcam, ab211033, 1:1000), FBXO33 (Novus Biologicals, NBP1-91890, 1:1000), Thrombospondin 1 (Santa Cruz, sc-393504, 1:500), Colla2 (Santa Cruz, sc-393573, 1:500), NRP2 (Santa Cruz, sc-13117, 1:500), FAK (Cell Signaling Technology, 3285, 1:1000), Phospho-FAK (Tyr397) (Cell Signaling Technology, 3283, 1:1000), Phospho-FAK (Tyr576/577) (Cell Signaling Technology, 3281, 1:1000), Phospho-FAK (Tyr925) (Cell Signaling Technology, 3284, 1:1000), RUNX2 (Abcam, ab23981, 1:1000), SP7 (Abcam, ab22552, 1:1000).

### Co-immunoprecipitation assays

For endogenous co-IP, BMSCs were treated with sciadopitysin or vehicle. We incubate total cell lysates overnight at 4°C with YBX1 antibody (4202, Cell Signaling Technology) or IgG as control. We used Dynabeads Protein G to collect the antigen-antibody complexes. After three times washes, the complexes were subjected to immunoblotting with appropriate antibodies.

For exogenous co-IP, BMSCs were transfected with a series of HA-tagged mutated YBX1 plasmid and His-tagged FBXO33 plasmid using Lip2000. Two days after transfection, total cell lysates were collected and incubated with antibody (ab9108; anti-His tag antibody; Abcam) overnight at 4°C. Dynabeads Protein G was used to collect the antigen-antibody complexes. After three times washes, the complexes were subjected to immunoblotting with appropriate antibodies.

### Molecular docking

Molecular docking was conducted as previously described ^27^. Briefly, the structure of mouse YBX1 was modeled on the basis of structure of human YBX1 (PDB code:1H95) using MODELER software for their high homology as previously described ^27^. We performed virtual screening between the natural small compounds library (Target Mol, US, Boston) and YBX1 through Autodock Vina and Dock 6.7.We used the autodock tools (ADT) to set the virtual screening parameters. A small number of top-ranked compouds were purchased from Target Molecule Corp and used as candidates for further study.

### Micro-CT analysis

The femurs were fixed in 4% paraformaldehyde for 24 hours, then scanned and analyzed by high-resolution µCT (VIVACT 80; SCANCO Medical AG, Switzerland). Scanner was set at a current of 145 μA and a voltage of 55 kV with a resolution of 15 μm per pixel. The image reconstruction software (NRecon, version 1.6, Bioz), data analysis software (CT Analyser, version 1.9, Bruker microCT) and 3-dimensional model visualization software (μCT Volume, version 2.0, Bruker microCT) were used to analyse the BV/TV, Tb. Th, Tb. N, and Tb. Sp of the distal femoral metaphyseal trabecular bone. The region of interest was defined as 5% of femoral length below the growth plate.

### Study population

Human bone marrow samples were obtained from patients undergoing knee joint replacement because of osteoarthritis, undergoing hip joint replacement because of femoral neck and/or femoral head fractures or undergoing open reduction internal fixation because of tibia or femur shaft fractures. Human bone marrow aspiration and collection were conducted by the Orthopedic Surgery Department at Xiangya Hospital of Central South University. A total of 60 patients (30 male and 30 female) were selected on the basis of the inclusion and exclusion criteria. All subjects were screened using a detailed questionnaire, disease history, and physical examination. Subjects were excluded from the study if they had conditions affecting bone metabolism or previous pathological fractures within 1 year or had received treatment with glucocorticoids, estrogens, thyroid hormone, parathyroid hormone, fluoride, bisphosphonate, calcitonin, thiazide diuretics, barbiturates, antiseizure medication.

### Statistics

The data are expressed as mean ± SEM. The data is normally distributed, Two-tailed Student’s t test is used to compare between two groups. One-way analysis of variance (ANOVA) is used to compare the difference between multiple groups. The statistics is applied by SPSS 20.0. Statistical differences were supposed to be significant when P<0.05.

## Author Contribution

Y.X designed the experiments and generated data; Y.X and GP.C carried out majority of the experiments; X.F, Q.G, T.S, Y.H and CJ.L contributed to acquiring data; XH.L, YJ.Z and M.Y co-advised the study; GP.C and M.Y wrote the manuscript; YJ.Z and M.Y is the guarantor of this work and has full access to all the data in this study and takes the responsibility for data accuracy.

## Acknowledgements

This work was supported by grants from National Natural Science Foundation of China (Grant No. 92149306, 82120108009, 81930022, 81900810, 82170903, 81900732, 82000811), the National Postdoctoral Program for Innovative Talents of China Postdoctoral Science Foundation (Grant No: BX20200390), China Postdoctoral Science Foundation (Grant No: 2021M703642) and Hunan Provincial Science and Technology Department (Grant No: 2020RC2011).

## Supplementary Materials: Including Figs. S1 to S5

**Supplementary figure 1.**
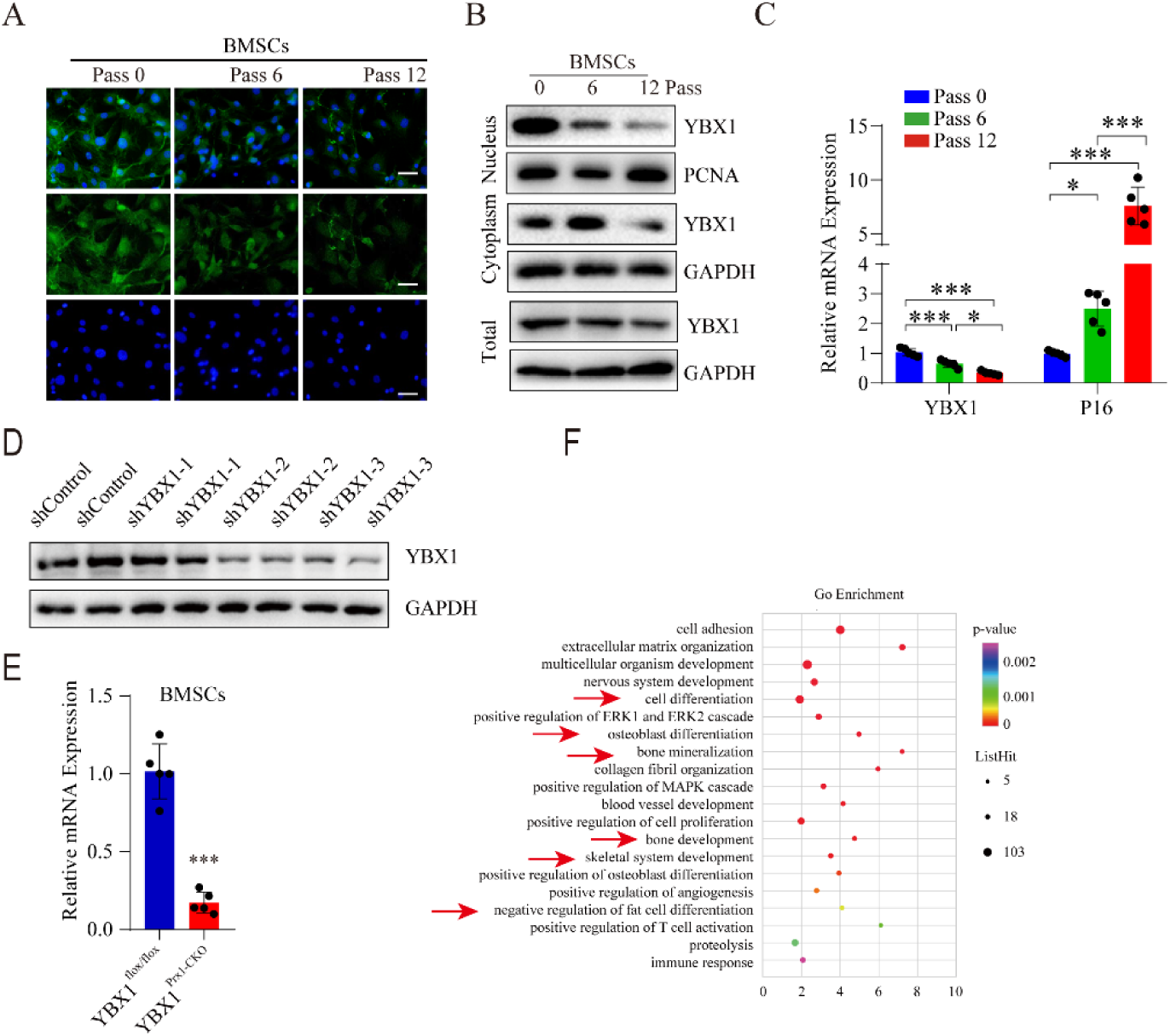
YBX1 level was lower in cultured primary BMSCs from late passage. (A) Representative images of immunofluorescence staining of YBX1 (green) in primary BMSCs isolated from passage 0, passage 6 and passage 12. Scale bar: 100μm. (B) Western blot analysis of the expression of YBX1 in BMSCs from passage 0, passage 6 and passage 12. (C) qRT-PCR analysis of the expression of *Ybx1* and *P16* in BMSCs from passage 0, passage 6 and passage 12. (D) Western blot analysis of the level of YBX1 in BMSCs transfected with adenovirus driven control and YBX1 shRNA. (E) qRT-PCR analysis of the levels of *Ybx1* in BMSCs isolated from *YBX1^Prx1-CKO^* mice and *YBX1^flox/flox^*mice. (F) GO analysis of differentially expressed genes in BMSCs isolated from *YBX1^Prx1-CKO^* mice and *YBX1^flox/flox^* mice. Data shown as mean ± SEM. *, *P* < 0.05; **, *P* < 0.01; ***, *P* < 0.001; One-way ANOVA for C; Student’s t test for E.

**Supplementary figure 2.**
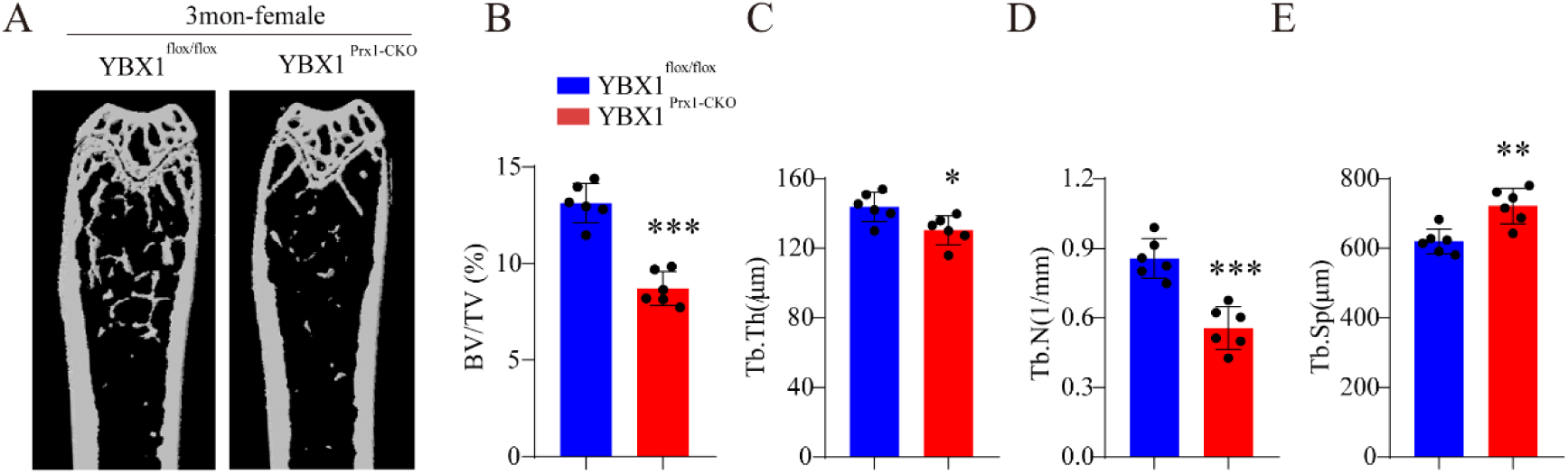
Depletion of YBX1 in BMSCs results in accelerated bone loss in female mice. (A-E) Representative μCT images (A) and quantitative μCT analysis of trabecular bone microarchitecture (B-E) in distal femora from 3-month-old female *YBX1^Prx1-CKO^*mice and *YBX1^flox/flox^* mice. Data shown as mean ± SEM. ^#^, no significant difference; *, *P* < 0.05; **, *P* < 0.01; ***, *P* < 0.001; Student’s t test.

**Supplementary figure 3.**
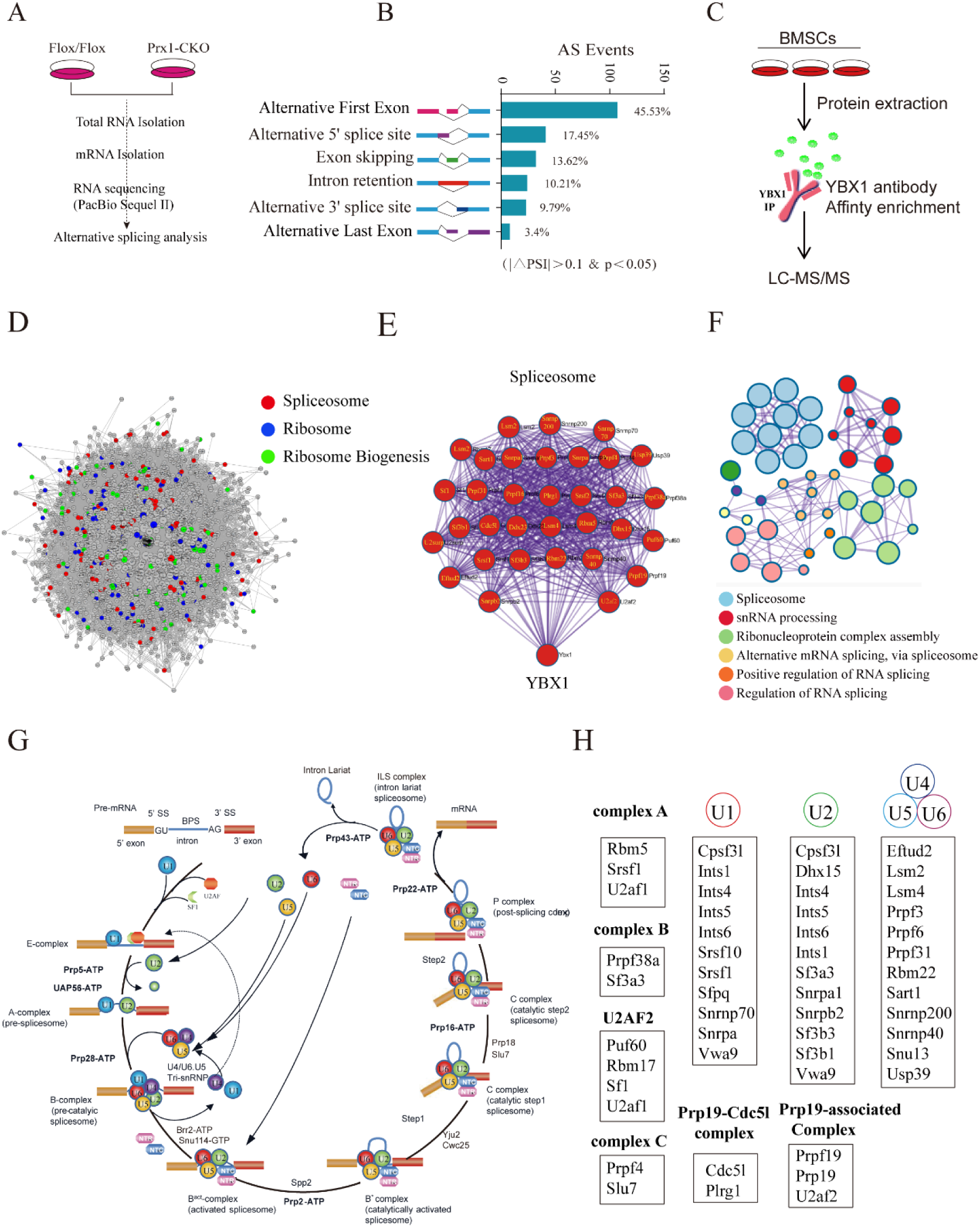
YBX1 interacted with spliceosome components and YBX1 deficiency altered pre-mRNA splicing in BMSCs. (A) Schematic diagram of experimental process of alternative splicing analysis. (B) Histogram of the differentially spliced events between BMSCs isolated from *YBX1^Prx1-CKO^* mice and *YBX1^flox/flox^* mice. (C) Study design of mouse YBX1 interactomes. YBX1 interactomes are analyzed by LC-MS/MS. (D) The network represents the proteins interacting with YBX1 in BMSCs. Specific color of the node indicates spliceosomal, ribosomal and ribosomal biogenesis proteins. (E) Network representation of YBX1 interacting spliceosomal proteins in BMSCs. (F) Enrichment network representing the top 10 enriched terms of YBX1 related proteins. Enriched terms with high similarity were clustered and rendered as a network, while each node represents an enriched term and is colored according to its cluster. Node size indicates the number of enriched proteins. (G) Spliceosome proteins interacting with YBX1 participate in spliceosome assembly reaction in a stepwise manner to form a mature mRNA. (H) Summarizing important YBX1 interacting spliceosomal proteins based on their spliceosome complex.

**Supplementary figure 4.**
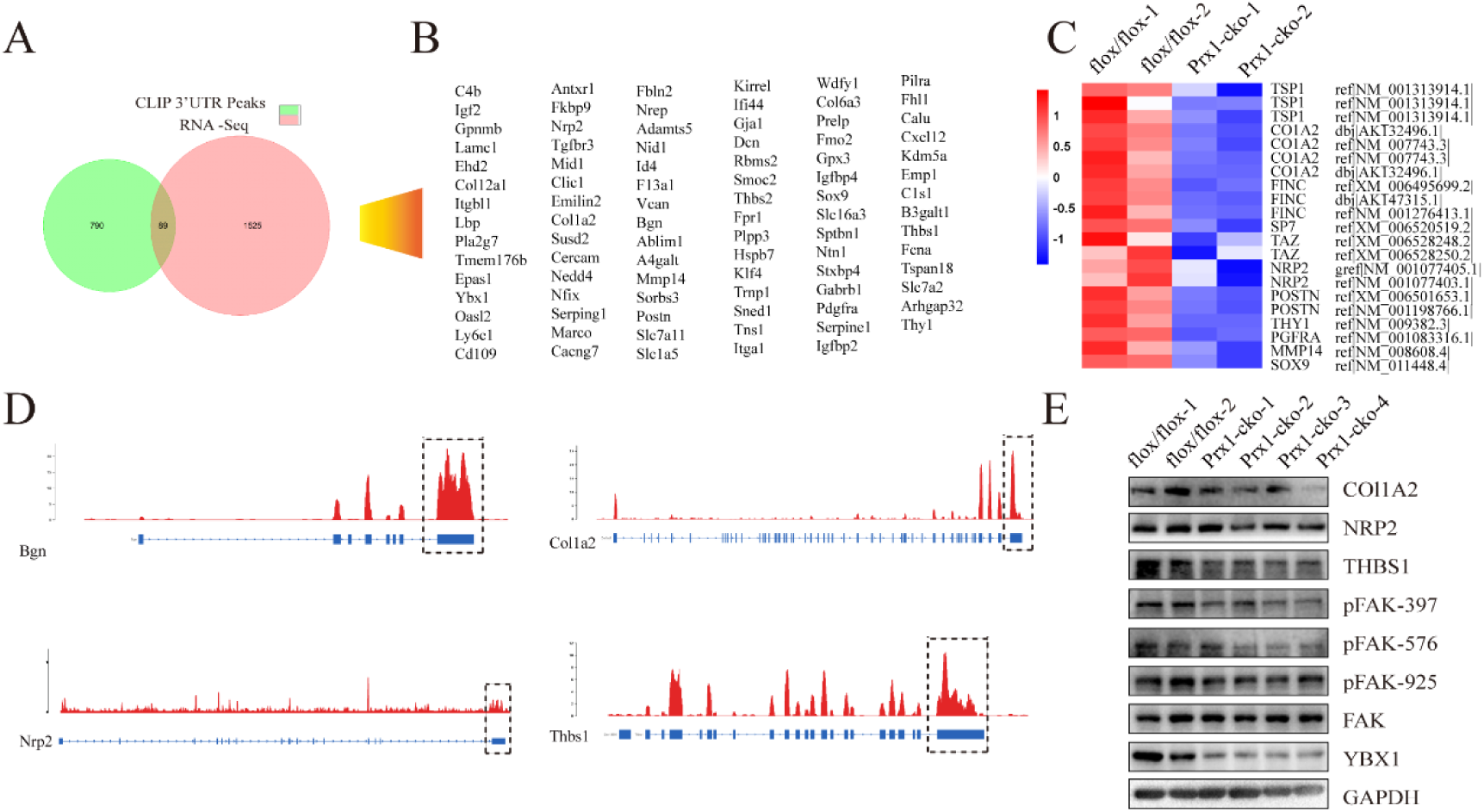
YBX1 alters mRNA stability by binding to 3’UTR. (A) Venn diagrams of overlapping genes targeted by YBX1 on 3’UTR are and showed altered expression upon YBX1 deletion. (B) Genes targeted by YBX1 on 3’UTR are and showed altered expression upon YBX1 deletion. (C) Heat map of differentially expressed genes in BMSCs isolated from *YBX1^Prx1-CKO^* mice and *YBX1^flox/flox^* mice. (D) RNA-seq read showed YBX1 binding the 3’UTR area of *Bgn, Colla2, Nrp2* and *Thbs1* mRNA in BMSCs. (E) Western blot analysis of the expression of YBX1, NRP2, THBS1, FAF and phosphorylation of FAK in BMSCs isolated from *YBX1^Prx1-CKO^* mice and *YBX1^flox/flox^*mice.

**Supplementary figure 5.**
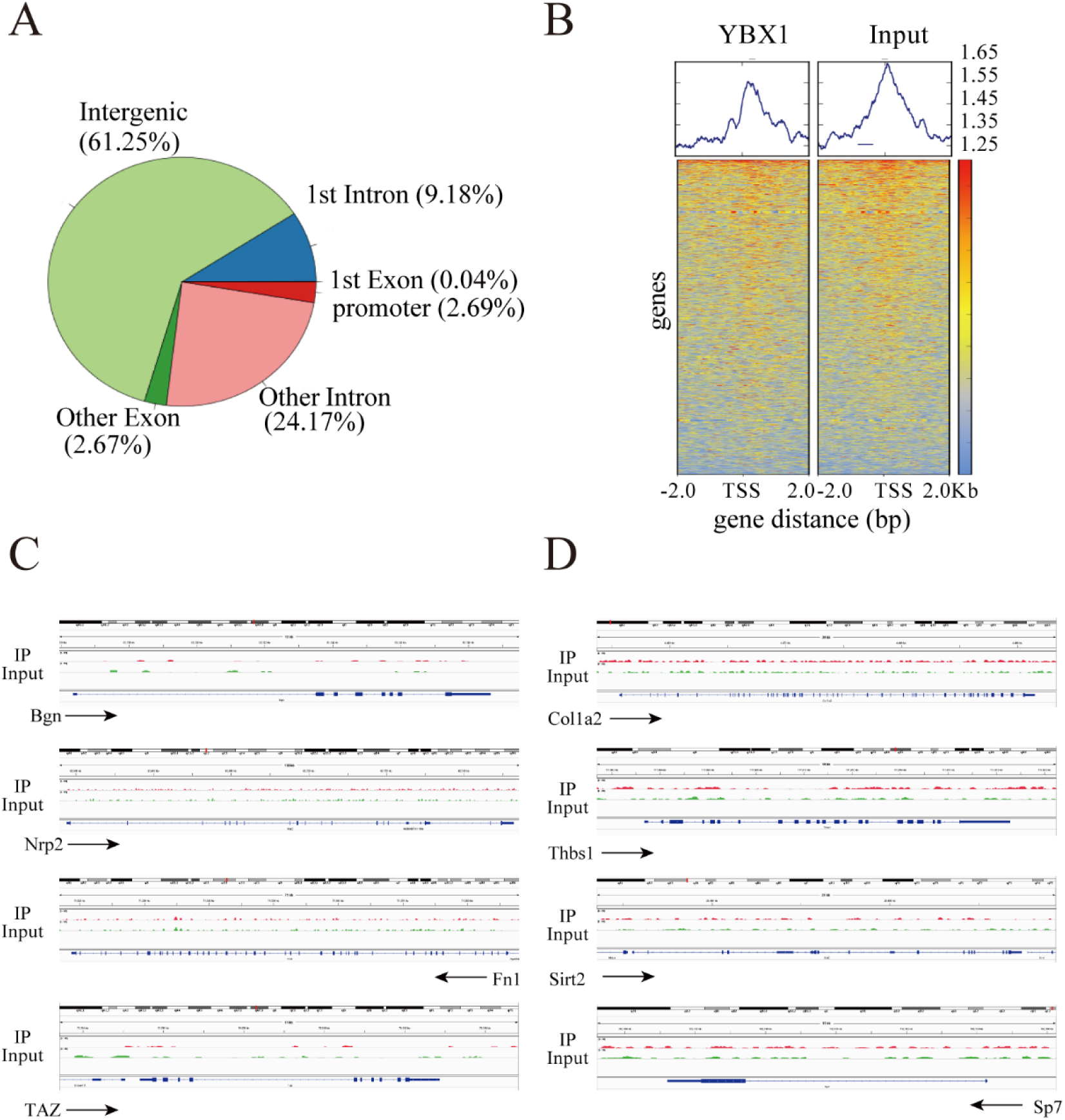
YBX1 do not bind to the promoter regions of *Bgn, Colla2, Nrp2, Thbs1, Fn1, Taz, Sirt2*, and *Sp7* in BMSCs. (A) Distribution of YBX1 ChIP-seq peaks in the genome. (B) Heatmap of YBX1 and imput ChIP-seq peaks in BMSCs. The signal is displayed within 2 kb of transcriptional start site. (C) ChIP-seq profile for YBX1 in BMSCs at *Bgn, Nrp2, Fn1* and *Taz*. (D) ChIP-seq profile for YBX1 in BMSCs at *Colla2, Thbs1, Sirt2*, and *Sp7*.

